# Nuclear speckles comprise hierarchical nanoscale interaction networks that shape molecular dynamics

**DOI:** 10.64898/2026.10.30.763187

**Authors:** Minxue Liu, Tianyou Yao, Jonas Zähringer, You Jin Song, Qingcai Meng, Daiki Futagi, Astrid D. Haase, Steven H DeVries, Susan M. Freier, Supriya G. Prasanth, Taekjip Ha, Ido Golding, Kannanganattu V. Prasanth

## Abstract

Biomolecular condensates organize essential cellular processes, but how their internal architecture regulates molecular behavior in living cells remains poorly understood. Here, we use MINFLUX nanoscopy for quantitative single-molecule imaging and live-cell tracking to resolve the organization and dynamics of specific macromolecules within nuclear speckles. Rather than behaving as uniform liquid-like compartments, nuclear speckles are hierarchically organized into dynamic nanoscale interaction networks. Proteins and RNAs assemble into molecular clusters that generate transient interaction nanodomains, where individual molecules alternate between diffusion and confinement while exploring higher-order networks. Physiological and genetic perturbations selectively remodel molecular clustering and nanodomain organization, and these changes are not consistently coupled to changes in the abundance of the measured components. Together, these findings reveal a multiscale organization of nuclear speckles in which molecular clustering, transient confinement, and dynamic exchange represent distinct but interconnected features of condensate architecture in living cells.

---

The formation of biomolecular condensates has emerged as a fundamental principle of cellular organization, whereby condensates concentrate proteins and nucleic acids to regulate diverse molecular processes, including transcription and RNA processing.^1–3^ Rather than being enclosed by membranes, biomolecular condensates assemble through multivalent interactions among proteins and RNAs, forming dynamic compartments whose components exchange continuously with the nucleoplasm or cytoplasm.^2^

Earlier studies emphasized biomolecular condensates as homogeneous dense phases formed through liquid-liquid phase separation (LLPS).^1–3^ However, computational and *in vitro* studies have shown that condensates can also contain spatially heterogenous, percolated interaction networks, in which interconnected molecular interactions form a network spanning the condensate.^1–12^ RNA binding proteins with RNA recognition motifs (RRMs) can form size-limited nanoscale assemblies, or microphases, that are distinct from the macroscopic condensates expected from canonical LLPS.^13–15^ Hierarchical organization, in which smaller molecular assemblies are spatially organized into larger structures, is predicted to generate distinct local environments that modulate encounter frequencies, reaction rates, and biochemical specificity. However, direct evidence for such multiscale organization within condensates in living cells has remained limited by the spatial and temporal resolution of conventional imaging.

Nuclear speckles provide an ideal model to test the hypotheses emerging primarily from *in vitro* reconstitutions. These nuclear condensates are enriched in RNA binding proteins and RNAs, including structural scaffolds (SON and SRRM2), splicing regulators (SRSF-family proteins), and noncoding RNAs (*MALAT1* and snRNAs), and are linked to transcription and pre-mRNA processing.^13, 16–19^ Their assembly is thought to arise through multivalent interactions among intrinsically disordered regions and RRMs,^13, 17–21^ while speckle proximity to actively transcribed genes indicates a functional role in coordinating RNA metabolism.^20, 22–40^ Nuclear speckles possess heterogeneous internal organization rather than behaving as homogeneous liquid compartments, ^22, 41, 42^ but how this heterogeneity is structured at the nanoscale remains unknown.^13, 17–21^

Reconstituted splicing factors such as SRSF1 undergo nanoscale microphase separation below the diffraction limit of conventional light microscopy.^13, 20^ Speckle-resident *MALAT1* promotes SRSF1 microphase assembly *in vitro*. ^13^ Furthermore, within the nucleus, *MALAT1* enhances SRSF1 recruitment to active transcription sites, facilitating pre-mRNA processing.^20^ These findings suggest that microphases modulate local concentrations of splicing regulators near active genes.^13, 20^ However, whether such nanoscale assemblies exist *in vivo*, how they constrain single-molecule diffusion, and whether they remodel during cellular stress remain unresolved.

Here, we combine MINFLUX nanoscopy^43, 44^ for quantitative single-molecule imaging and live-cell tracking to resolve the internal architecture and dynamics of nuclear speckles at nanometer spatial precision and sub-millisecond temporal resolution.^45^ We show that in living cells, nuclear speckles are hierarchically organized into nanoscale interaction environments that govern molecular mobility. Furthermore, these interaction landscapes are selectively remodeled by perturbations to transcription, RNA splicing, stress signaling, electrostatic interactions, and scaffold composition. Together, our results directly link nanoscale spatial clustering to transient molecular confinement in living cells, demonstrating that internal condensate architecture can be dynamically regulated independently of gross molecular abundance.

## Results

### Quantitative mapping reveals the molecular inventory of nuclear speckles

To determine the molecular composition and spatial organization of nuclear speckles at nanometer resolution, we established a single-molecule MINFLUX imaging framework for quantitative visualization of speckle-enriched proteins and RNAs. This approach enabled direct localization and counting of individual molecules inside and outside nuclear speckles. MCF7 cell lines carrying HaloTags in the endogenous gene loci of three speckle-enriched splicing proteins SRRM2, SON, and SRSF1 were generated to produce N-terminal fusions, followed by labeling with a fluorescent HaloTag ligand (Fig.1a and Extended Data Fig.1a, see Methods).^46^ Unlike conventional immunofluorescence labeling,^47^ HaloTag labeling provides a defined 1:1 labeling stoichiometry between HaloTag fusion proteins and fluorophores, facilitating molecular counting. As a reference sample, separate MCF7 cells were transfected with a plasmid expressing HaloTag alone and imaged under the same conditions (Extended Data Fig.1b). The speckle-enriched ncRNAs *U2 snRNA* and *MALAT1* were detected by single-molecule fluorescence in situ hybridization (smFISH) using single ∼45-nt sulfo-Cy5 (sCy5)-conjugated oligonucleotide probes targeting *U2 snRNA* or the 3’ region of *MALAT1* (Fig.1b and Extended Data Fig.1a).^48, 49^ To assess whether an individual smFISH probe could be detected by MINFLUX, we examined two reference conditions expected to lack specific RNA targeting: smFISH in MCF7 using probes targeting PP7, a non-mammalian RNA originally from bacteriophage^50^ and smFISH using a probe against *MALAT1* in *MALAT1*-knockout MCF7 cells.^20^ Under both conditions, MINFLUX imaging revealed sparse and spatially isolated fluorescent spots (Extended Data Fig.1c–d). These results support the use of single-probe labeling for MINFLUX imaging, and provide a defined 1:1 correspondence between fluorophore and RNA molecule, unlike conventional multi-probe strategies,^51^ thereby facilitating accurate RNA quantification.

**Fig. 1:**
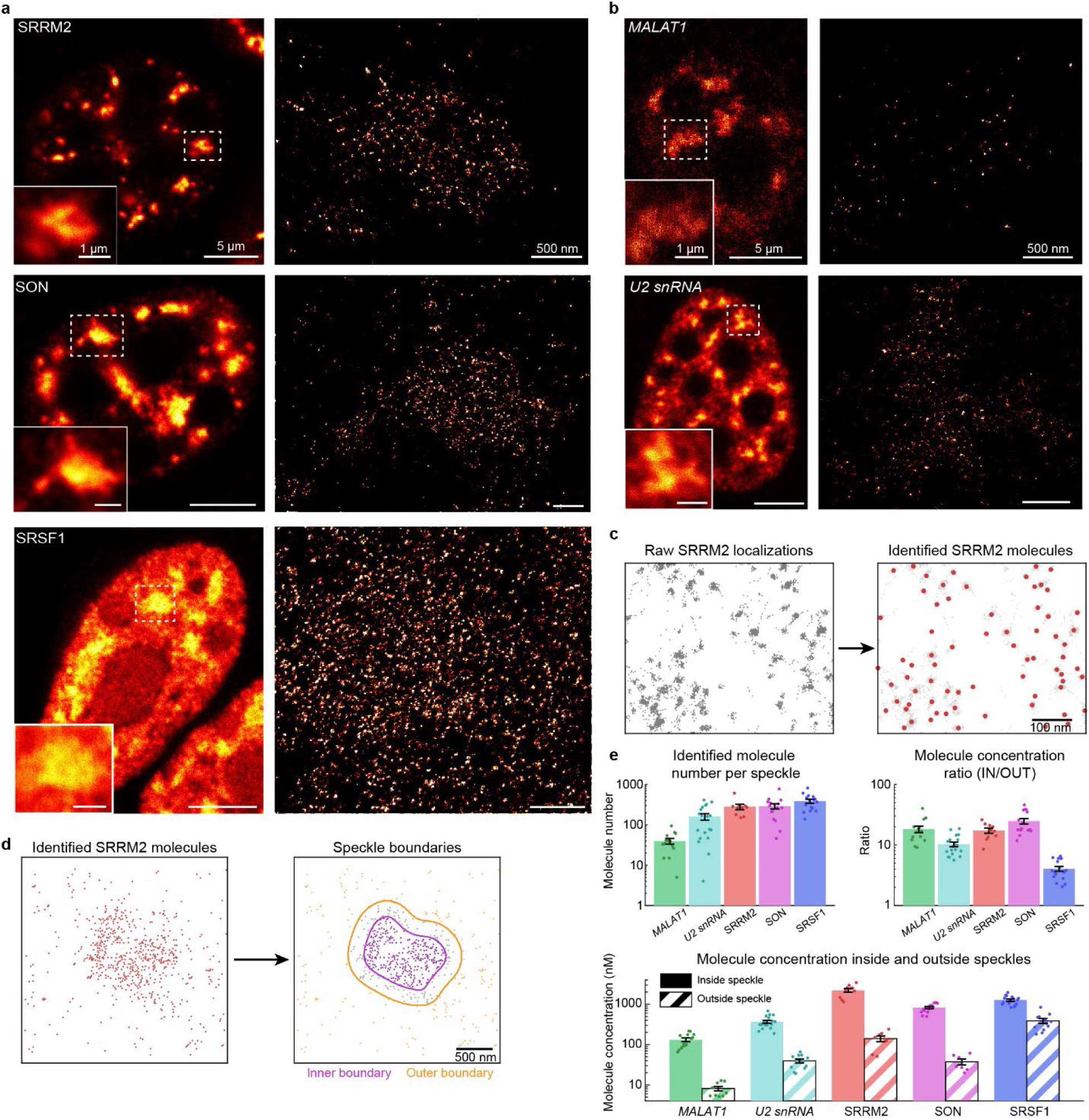
MINFLUX imaging and quantification of individual speckle proteins and RNAs in fixed cells. **a,** Representative 2D MINFLUX images of SRRM2 (top), SON (middle), and SRSF1 (bottom). Left, confocal images acquired on the MINFLUX microscope showing the nucleus. Inset, magnified view of a representative nuclear speckle used for MINFLUX imaging. Right, 2D MINFLUX images of the corresponding nuclear speckles. Scale bars, 5 μm (left), 1 μm (inset), and 500 nm (right). **b,** Representative 2D MINFLUX images of *MALAT1* (top) and *U2 snRNA* (bottom). Left, confocal images acquired on the MINFLUX microscope showing the nucleus. Inset, magnified view of a representative nuclear speckle used for 2D MINFLUX imaging (right). Scale bars, 5 μm (left), 1 μm (inset), and 500 nm (right). **c,** Identification of individual SRRM2 molecules from MINFLUX localizations. Left, raw MINFLUX localizations. Right, identified molecule positions (red) overlaid on the corresponding MINFLUX localizations (gray). **d,** Determination of nuclear speckle boundaries. Left, identified SRRM2 molecules. Right, inner and outer speckle boundaries. **e,** Quantification of protein and RNA abundance inside and outside nuclear speckles. Top left, number of identified molecules per speckle. Top right, ratio of molecular concentrations inside to outside speckles. Bottom, molecular concentrations inside and outside speckles. Data are presented as mean ± SEM. From left to right, n = 14, 19, 10, 16 and 16 inside speckles, and n = 10, 12, 8, 7 and 12 outside speckles.

To extract molecular positions from the raw MINFLUX data, we assessed whether the trace identifiers (TIDs) provided in MINFLUX datasets could be used to identify individual molecules. In MINFLUX datasets, localizations assigned to the same fluorescence emission event are grouped under a common TID,^47, 52, 53^ and TIDs have been proposed as molecular identifiers.^47, 53^ However, our data challenged this interpretation: the nearest-neighbor distance (NNd) distributions between trace positions exhibited a pronounced peak at ∼3nm (Extended Data Fig.1e). This peak was also observed in the control datasets (HaloTag-alone, PP7-targeting probes, and *MALAT1*-knockout cells) (Extended Data Fig.1e), where signals are expected to arise predominantly from isolated fluorophores, implying repeated detections of the same molecule rather than neighboring molecules. This behavior was also evident visually in a representative SRRM2 dataset, in which trace centers were closely spaced and localizations assigned to different TIDs frequently overlapped in space (Extended Data Fig.1f). Such observations are consistent with previous reports that a single fluorophore may give rise to multiple temporally separated localization events,^54, 55^ rendering trace-based counting unreliable in our data analysis.

Instead of relying on trace identifiers, we adopted a localization-based strategy for molecule identification (Extended Data Fig.1g, see Methods). Briefly, traces containing fewer than three localizations were discarded, consistent with previous MINFLUX analyses.^56–58^ The remaining localizations were then clustered using the Density-Based Spatial Clustering of Applications with Noise (DBSCAN) algorithm.^57, 59^ To resolve clusters that could contain localizations from multiple nearby molecules, we further applied a Gaussian mixture model (GMM)-based decomposition procedure,^59, 60^ which partitioned such clusters into multiple subclusters whose localization coordinates were modeled by two-dimensional Gaussian components. The resulting subclusters, together with DBSCAN clusters that did not require further subdivision, were operationally treated as individual-molecule localization aggregates (Fig.1c and Extended Data Fig.1h). This interpretation was supported by multiple observations. First, across all examined species, the distributions of the radii of gyration of localizations assigned to these identified molecules exhibited a common peak at ∼6 nm (Extended Data Fig.1i). The same molecule identification pipeline applied to the control datasets yielded a comparable peak (Extended Data Fig.1i). The position of this peak was compatible with the finite spatial spread expected from repeated localizations of single fluorophores under MINFLUX imaging.^43, 44^ In addition, diffraction-limited images reconstructed from the inferred molecular positions closely resembled the corresponding confocal images acquired from the same regions (Extended Data Fig.1j), indicating that the inferred molecular positions preserve the overall spatial distribution of fluorophores.

With molecular positions determined, we next quantified the abundance of speckle-enriched proteins and RNAs localized inside and outside nuclear speckles. To delineate speckle regions based on the discrete molecular positions, we reconstructed a continuous molecular density field as diffraction-limited images from the inferred molecule positions and defined inner and outer speckle boundaries based on the resulting molecular density profiles, which exhibited an approximately constant interior plateau followed by a gradual decrease at the speckle periphery (Extended Data Fig.1j; see Methods). These boundaries were subsequently used to define regions inside and outside speckles for quantitative analyses (Fig.1d and Extended Data Fig.1j). Quantification within the imaged speckle regions revealed ∼300-500 molecules of SRRM2, SON, and SRSF1, together with ∼200 *U2 snRNA* and ∼50 *MALAT1* RNA (Fig.1e).

We next estimated molecular concentrations inside and outside speckles and determined the enrichment inside speckles relative to the nucleoplasm. Two-dimensional molecular densities were converted to volumetric concentrations assuming a finite axial sampling depth (see Methods). Concentration estimates for HaloTag-labeled proteins were corrected for knock-in efficiencies measured by Droplet Digital PCR (SRRM2, SON) and western blot (SRSF1) (Extended Data Fig.1k–l). The scaffolding proteins SON and SRRM2 exhibited concentrations of ∼800 nM and ∼2.2 µM, respectively. The abundance of SRRM2 observed here is broadly consistent with previous estimates from fluorescence-based measurement and mass spectrometry-based measurements.^18, 61^ SRSF1 concentration was ∼1.3 µM, whereas *U2 snRNA* and *MALAT1* RNA were present at ∼400 nM and ∼150 nM, respectively (Fig.1e). Most speckle components exhibited >10-fold enrichment in the speckle interior, except for SRSF1 which displayed only ∼4-fold enrichment (Fig.1e). These results establish a quantitative single-molecule quantification of nuclear speckle components.

### Intermolecular clustering establishes the nanoscale architecture of nuclear speckles

Single-molecule localization of SRRM2 suggested a spatially heterogeneous distribution of molecules within individual speckles (Fig.2a), and similar spatial patterns were observed across all examined proteins and RNAs (Extended Data Fig.1h). To test whether these density fluctuations reflect genuine molecular clustering clearly distinguishable from complete spatial randomness (CSR: a spatial Poisson process),^62^ we analyzed the nearest-neighbor distance distributions (Fig.2b and Extended Data Fig.2a) and pair correlation function g(r)^63, 64^ (Fig.2c and Extended Data Fig.2b) compared with CSR models at matched concentrations.

**Fig. 2:**
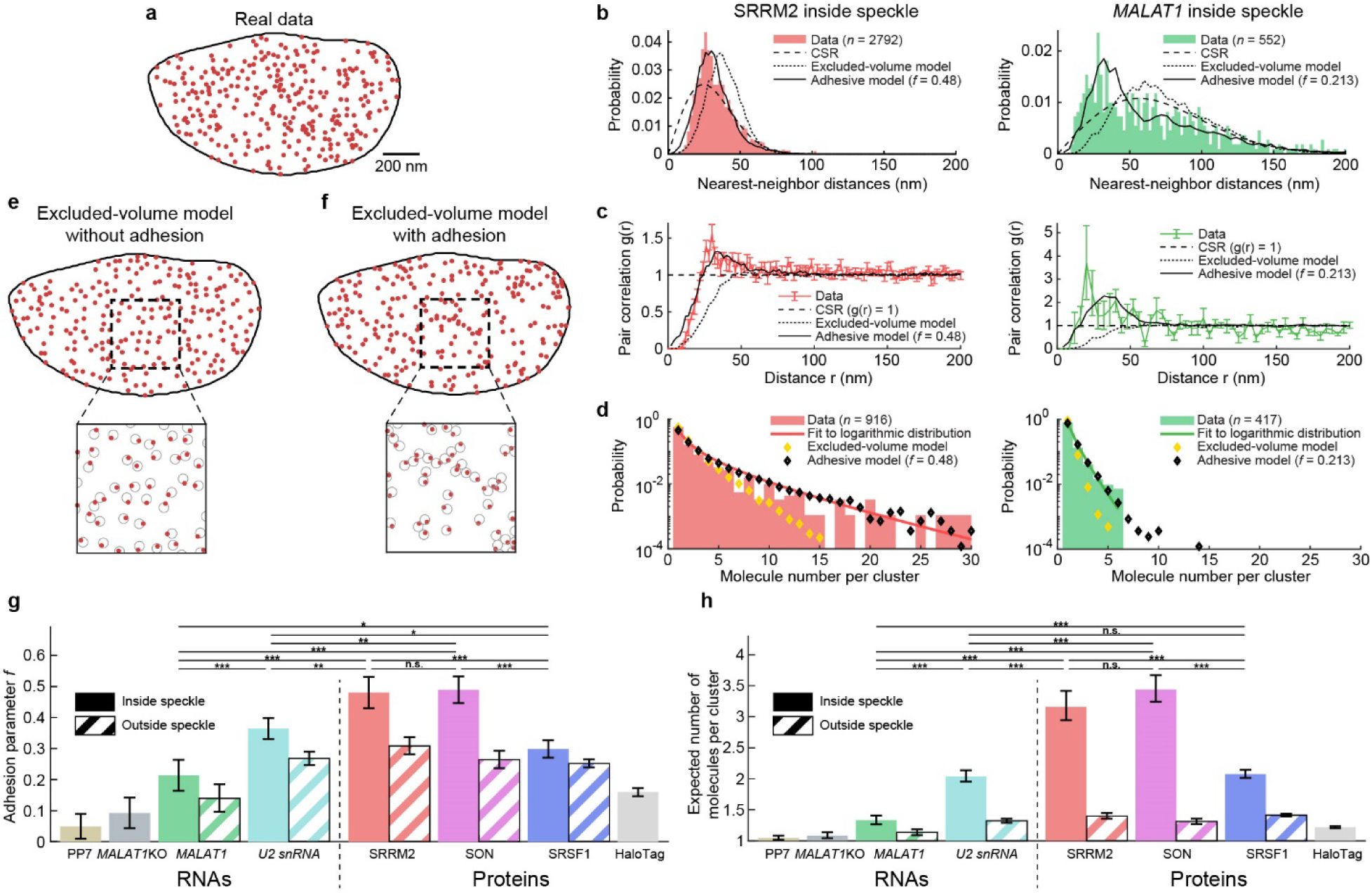
Speckle components form molecular clusters inside nuclear speckles. **a,** Representative spatial distribution of SRRM2 molecules within a nuclear speckle. **b,** Nearest-neighbor distance (NNd) distributions of SRRM2 (left) and *MALAT1* (right) inside nuclear speckles, pooled across speckles (bars), compared with complete spatial randomness (CSR; dashed lines), the excluded-volume model without local molecular adhesion (dotted lines), and the excluded-volume model with local molecular adhesion (solid lines). **c,** Pair correlation functions, g(r), of SRRM2 (left) and *MALAT1* (right) molecules inside nuclear speckles, averaged across speckles (colored lines with error bars), compared with CSR (dashed lines), the excluded-volume model without local molecular adhesion (dotted lines), and the excluded-volume model with local molecular adhesion (solid black lines). **d,** Distributions of the number of molecules per cluster for SRRM2 (left) and *MALAT1* (right) inside nuclear speckles, pooled across speckles (bars) and fitted to a logarithmic distribution (solid lines), compared with simulations from the excluded-volume model without local molecular adhesion (yellow markers) and the excluded-volume model with local molecular adhesion (black markers). **e,** Simulation of the excluded-volume model without local molecular adhesion. Top, simulated molecular distribution coordinates at the concentration measured inside nuclear speckles. Bottom, magnified view of the boxed region showing the excluded-volume disks (circles) and the simulated molecular coordinates (points) sampled within them. **f,** Simulations of the excluded-volume model with local molecular adhesion. Top, simulated molecular coordinates at the concentration and adhesion parameter *f* estimated for molecules inside nuclear speckles. Bottom, magnified view of the boxed region showing the excluded-volume disks (circles) and the simulated molecular coordinates (points) sampled within them. **g,** Adhesion parameter *f* for the five speckle components inside and outside nuclear speckles and in control datasets (HaloTag-alone, PP7-targeting probes, and *MALAT1*-knockout cells). **h,** The expected number of molecules per cluster for the five speckle components inside and outside nuclear speckles and in control datasets (HaloTag-alone, PP7-targeting probes, and *MALAT1*-knockout cells). Bar graphs in (**g**) and (**h**) show means with 95% confidence intervals (CIs). Asterisks denote non-overlapping CIs: (*) 95%, (**) 99%, (***) 99.9%; n.s. denotes overlapping 95% CIs.

Experimental NNd distributions and pair correlation functions deviated substantially from CSR expectations (Fig.2b–c and Extended Data Fig.2a–b). At distances < 15 nm, molecular pairs were less frequent, consistent with exclusion volume interactions. At intermediate distances (∼20-45 nm), NNd distributions and g(r) curves exhibited a pronounced peak across all species, with the peak position remaining similar despite differences in molecular concentration (Fig.2b–c and Extended Data Fig.2a–b). Similar deviations from CSR were also observed in the NNd distributions of molecules outside nuclear speckles (Extended Data Fig.2c). In contrast, the NNd distribution of the HaloTag-alone control, which lacks specific molecular interactions, showed closer agreement with CSR (Extended Data Fig.2d), further supporting local clustering of speckle components.

To quantify the size and composition of molecule clusters, we grouped the inferred molecular positions using DBSCAN (Extended Data Fig.1g), using a neighborhood radius (*ε* = 45 nm) corresponding to the characteristic intermolecular distance scale (∼20-45 nm) identified from NNd distributions and g(r) curves. Cluster-size distributions were dominated by small clusters but featured extended tails toward larger sizes (Fig.2d and Extended Data Fig.2e). These distributions were well fit by a logarithmic distribution (Fig.2d and Extended Data Fig.2e; see Methods).^65^ In contrast, speckle components localized outside of speckles, and negative controls (HaloTag-alone, PP7 probes, and *MALAT1*-knockout) showed little evidence of molecular clustering, with cluster-size distributions more strongly dominated by single-molecule clusters (Extended Data Fig.2f).

The reduced frequency of molecular pairs at short distances (Fig.2b–c and Extended Data Fig.2a–b) suggested a contribution from excluded-volume interactions. We therefore used a two-dimensional hard-disk model to test whether steric exclusion alone could account for the observed spatial organization. The disk radius was set to 15 nm, corresponding to the effective exclusion range inferred from the NNd distributions and pair correlation functions (Fig.2e; see Methods). At experimentally measured molecular concentrations, this model failed to reproduce the experimental NNd distributions and g(r) curves, and yielded substantially fewer large molecular clusters than observed experimentally (Fig.2b–d and Extended Data Fig.2a–b,e). Thus, steric exclusion alone is insufficient to account for the observed spatial organization.

We therefore extended the model by introducing an adhesion parameter *f*, representing an additional tendency for local inter-molecular adhesion beyond steric exclusion (Fig.2f). For each molecular species and condition, *f* was inferred from the measured molecular concentration and the fitted logarithmic-distribution parameter using an empirical mapping established from simulations (Extended Data Fig.2g; see Methods). Incorporating this additional adhesion recapitulated the experimental NNd distributions, g(r) curves, as well as the cluster size distributions, including the abundance of larger molecular clusters (Fig.2b–d and Extended Data Fig.2a–b,e).

Inside the speckles, the inferred adhesion parameter *f* was highest for the structural scaffolds SON and SRRM2, intermediate for SRSF1 and *U2 snRNA*, and lowest for *MALAT1* (Fig.2g). Molecules inside the nuclear speckles consistently exhibited higher *f* values than molecules located outside speckles or those in the control datasets (HaloTag-alone, PP7-targeting probes, and *MALAT1*-knockout cells), indicating a higher degree of local molecular adhesion inside the speckle environment (Fig.2g). For scaffolding proteins SRRM2 and SON, the inferred clusters contained on average ∼3 molecules, whereas SRSF1 and RNA displayed lower numbers per cluster (Fig.2h). Using a simple geometric approximation in which clusters were treated as compact arrangements of molecules with an exclusion radius of ∼15 nm, the observed average numbers of molecules per cluster correspond to estimated cluster diameters of ∼40-50 nm. Together, these results support a model in which enhanced local molecular adhesion within nuclear speckles contributes to the nanoscale organization of speckle components.

### Dynamic interaction environments shape molecular mobility in nuclear speckles

Although our analyses revealed heterogeneous molecular organization within nuclear speckles, how individual speckle components dynamically experience these local environments remains unclear. To directly probe nanoscale interactions and determine whether nanoscale molecular clustering (Fig.2f) affects molecular mobility, we tracked individual speckle proteins (SRRM2, SON and SRSF1) inside and outside of the speckles in living cells using live cell 2D MINFLUX tracking. The high spatial and temporal resolution of MINFLUX provides densely sampled single-molecule trajectories, enabling detailed characterization of molecular dynamics within individual trajectories.^44^ Endogenously expressed Halo-tagged SRRM2, SON, and SRSF1 were sparsely labeled with JFX650 and tracked at the single-molecule level (inside and outside speckles) in live MCF7 cells, with nuclear speckles marked by GFP-U2AF65 (Fig.3a). Individual protein molecules were tracked with a temporal resolution of ∼400 µs and a spatial precision of ∼16 nm for up to 4 s, during which they exhibited lateral displacements of up to several hundred nanometers, demonstrating substantial molecular mobility both inside and outside nuclear speckles (Fig.3b–c and Extended Data Fig.3a–d).

**Fig. 3:**
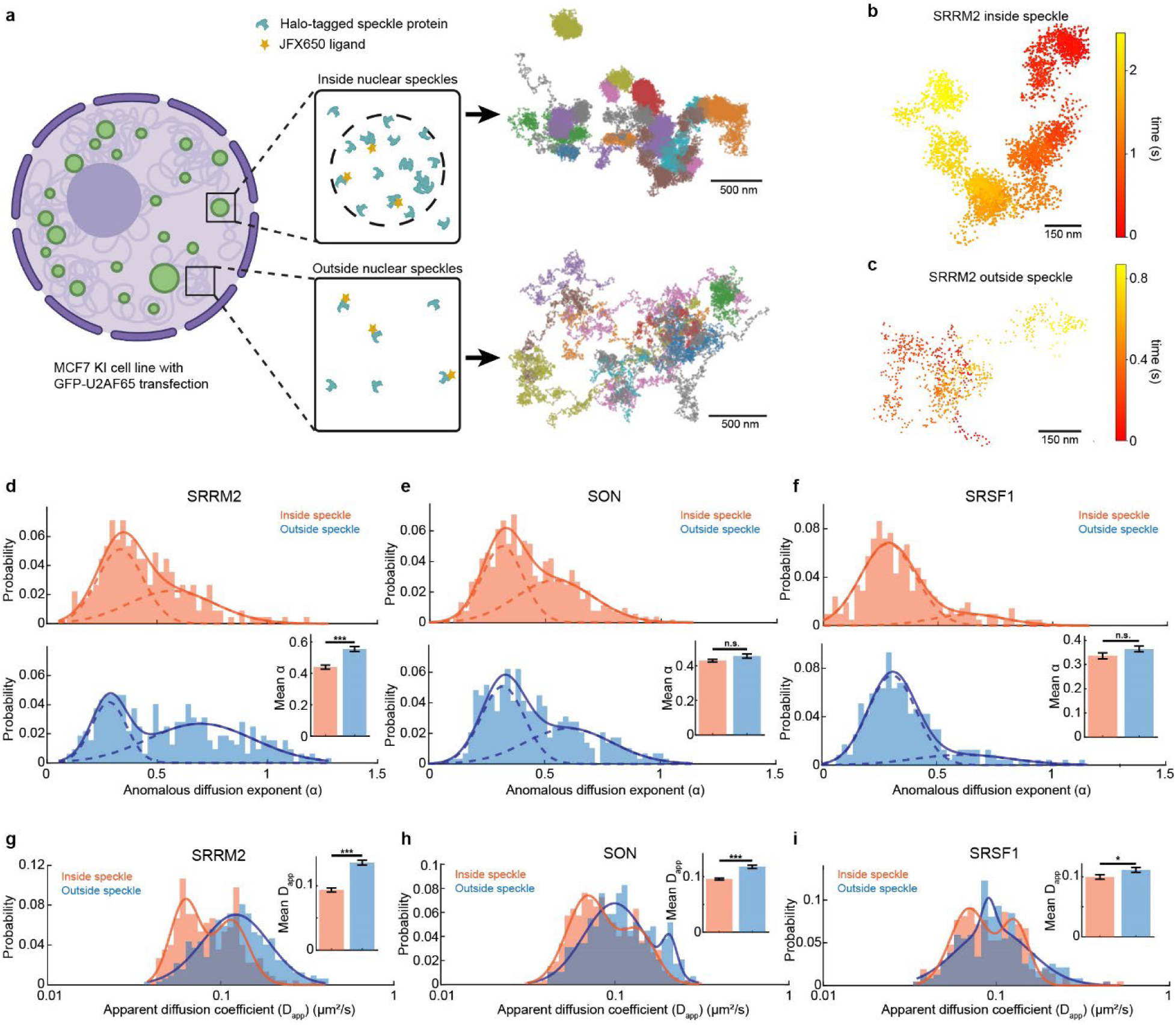
MINFLUX live tracking reveals multi-state diffusion behaviors of speckle proteins. **a,** Experimental design for MINFLUX tracking of speckle proteins. Halo-tagged SRRM2, SON, or SRSF1 was sparsely labeled with JFX650 HaloTag ligand in genome-edited MCF7 cell lines. GFP-U2AF65 was transiently expressed to mark nuclear speckles, and individual protein molecules were tracked by MINFLUX microscopy. Left, schematic of an MCF7 cell with nuclear speckles labeled by GFP-U2AF65 (green) and regions of interest (ROIs) selected inside and outside nuclear speckles. Right, all single-molecule trajectories of SRRM2 detected within a representative ROI inside (top) and outside (bottom) nuclear speckles, with individual trajectories shown in distinct colors. **b–c,** A representative single-molecule trajectory of SRRM2 inside (**b**) and outside (**c**) of nuclear speckles. Colors indicate the tracking time. **d–f,** Distributions of the anomalous diffusion exponent (α) for SRRM2 (**d**), SON (**e**), and SRSF1 (**f**) inside (top, orange) and outside (bottom, blue) nuclear speckles. α values are obtained by fitting the mean squared displacement (MSD) of individual trajectories to a power-law model. The distributions are fitted to a sum of two Gaussians as a guide to the eye; dashed lines show the individual Gaussian components and solid lines show their sum. Inset, mean α values for splicing factors inside and outside speckles. **g–i,** Distributions of apparent diffusion coefficients (D_app_) for SRRM2 (**g**), SON (**h**), and SRSF1 (**i**) inside (orange) and outside (blue) nuclear speckles. D_app_ values are estimated from MSD analysis of individual trajectories. The distributions are fitted to a sum of two Gaussians as a guide to the eye (solid lines). Insets represent mean D_app_ values for each protein inside and outside speckles. For **d–i**, α and D_app_ were obtained from the same trajectories: n = 225 and 321 for SRRM2, 524 and 290 for SON, and 209 and 236 for SRSF1, inside and outside speckles, respectively. Inset data are presented as mean ± SEM. (*) p ≤ 0.05, (**) p ≤ 0.01, and (***) p ≤ 0.001 (Mann–Whitney U test).

Since all tested speckle proteins exhibited higher degree of local molecular adhesion inside speckles (Fig.2g), we expected reduced molecular mobility inside speckles. We therefore quantified molecular motion by fitting the mean squared displacement (MSD) as a function of lag time (*τ*) to an anomalous 2D diffusion model, *MSD*(*τ*) = 4*Dτ^α^*, ^66, 67^ where D represents the generalized diffusion coefficient and α is the anomalous diffusion exponent (α=1: normal diffusion; α<1: sub diffusion) (Extended Data Fig.3e). The resulting α and D distributions spanned a broad range and displayed an apparently multi-component structure, indicating heterogeneous diffusive behaviors both inside and outside speckles among all tested speckle proteins (Fig.3d–f and Extended Data Fig.3g). Over 70% of the captured speckle proteins, whether inside or outside of the speckles, exhibited confined movement (α<0.7) (Extended Data Fig.3f).^68^ The broad distributions of D and α reflect diverse diffusive behaviors of speckle proteins and indicate the heterogeneous local environments inside speckles.

Next, we compared the diffusive behaviors of speckle proteins between inside and outside nuclear speckles through the calculation of the apparent diffusion coefficient (D_app_) from the short-time displacements of individual trajectories (see Methods). All three speckle proteins showed increased mobility outside speckles, as indicated by higher D_app_ (Fig.3–i). In contrast, *α* was higher outside speckles for SRRM2 but remained comparable between inside and outside speckles for SON and SRSF1 (Fig.3d–i). Overall, these results indicate that nuclear speckles provide a more constrained environment compared to the surrounding nucleoplasm.

Widefield single-particle tracking (SPT) was used to validate the MINFLUX tracking performance (Extended Data Fig.3h). Consistent with MINFLUX, widefield SPT showed substantially higher SRRM2 mobility outside speckles, and SON exhibited only a modest increase. While widefield SPT revealed a fast-diffusing SRSF1 population likely exceeding the MINFLUX tracking limit,^45^ a substantial slow-diffusing fraction of SRSF1 remained trackable (Extended Data Fig.3h).

Nuclear speckles are chromatin-depleted regions.^69^ Comparing speckle protein dynamics to chromatin-bound histone H2B (a benchmark for highly constrained, sub-diffusive motion^70^) showed that SRRM2 and SON exhibited significantly higher α values than H2B (Extended Data Fig.3a,i). Thus, while constrained, the speckle interior remains substantially more permissive to diffusion than the surrounding chromatin network.

### Speckle proteins dynamically explore hierarchically organized nanodomains

Visual inspection of MINFLUX tracking of the motion of speckle proteins at sub-millisecond timescales suggested that speckle proteins frequently switched between distinct diffusive states, repeatedly slowing within nanoscale regions (Fig.3b–c; Extended Data Fig.3a; Supplementary Videos 1–6). Such behavior is likely driven by transient local interactions and/or the heterogeneous molecular crowding^71, 72^ within nuclear speckles. To better characterize these multi-state diffusion behaviors, we segmented the trajectories based on the number of localizations and fitted the MSD of each segment using the power-law model described above (see Methods). Segments with *α* < 0.7 were classified as confined and those with *α* > 0.7 as diffusive (Fig.4a, see Methods).^68^ We fitted the MSD of the confined segments as a function of the lag time to a saturating exponential model (confined model)^73^ to estimate an effective radius of confinement (R_c_) (Fig.4b, see Methods). Inside speckles, all three proteins exhibited confinement within regions with a mean confinement radius R_c_ of ∼40 nm (Fig.4c and Extended Data Fig.4a–b). These confined regions are hereafter referred to as “speckle nanodomains”. Similar tracking in paraformaldehyde (PFA)-fixed cells produced substantially smaller confinement radii, primarily reflecting localization uncertainty and supporting that the larger confinement radii observed in live cells arise from genuine biological molecular dynamics within speckle nanodomains (Extended Data Fig.4c).

**Fig. 4:**
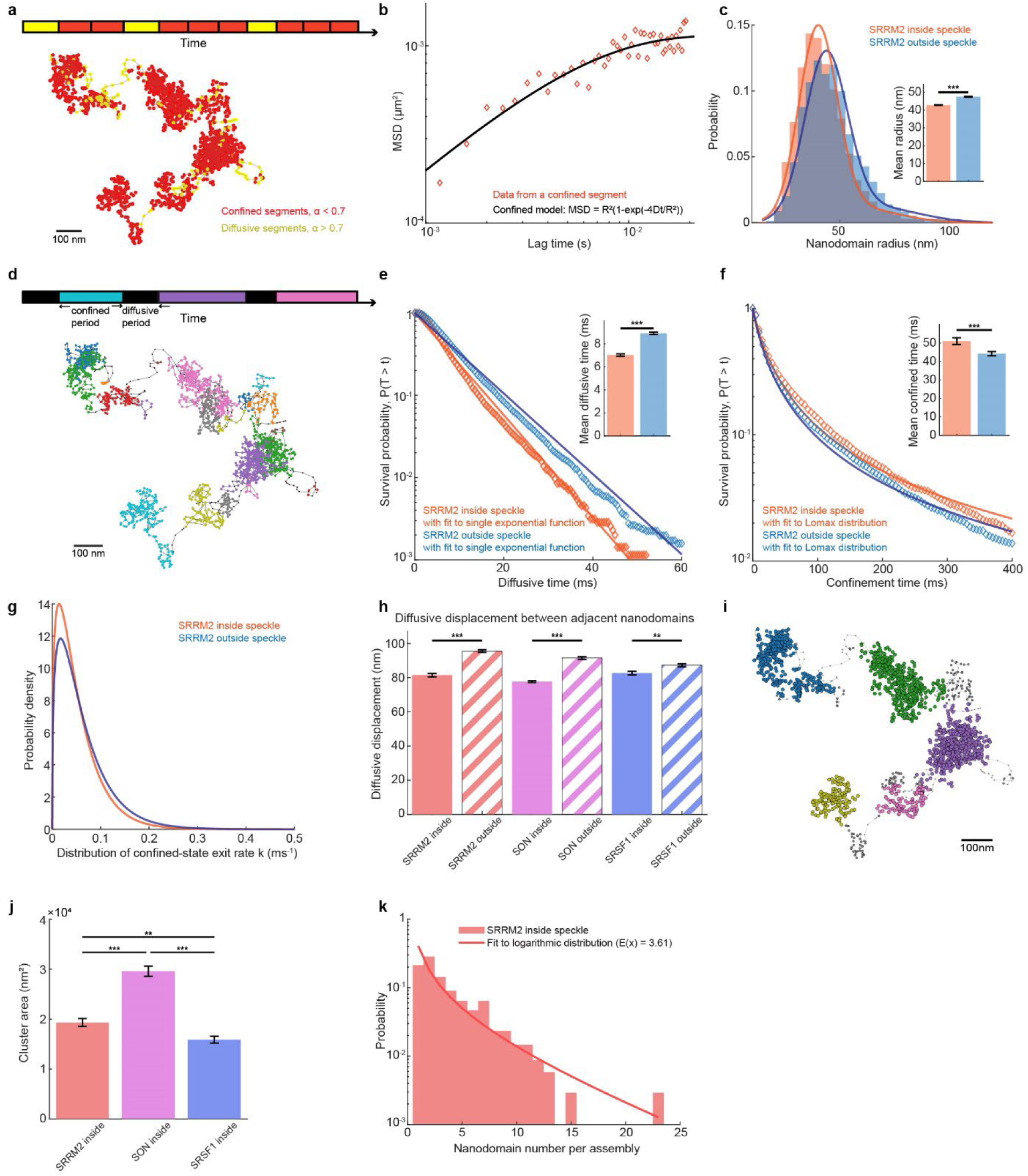
Speckle proteins transiently slow down within confined nanodomains. **a,** Identification of confined and diffusive segments within single-molecule trajectories. Individual trajectories are divided into segments, and the mean squared displacement (MSD) of each segment is fitted with a power-law model to determine the anomalous diffusion exponent α. Top, schematic showing the identified confined (red) and diffusive (yellow) segments over time. Bottom, a representative SRRM2 trajectory inside a nuclear speckle, with the corresponding confined and diffusive segments shown in red and yellow, respectively. **b,** MSD as a function of lag time for a representative confined segment of a SRRM2 trajectory inside a nuclear speckle (blue markers), with the fit to the confined model (red). **c,** Distributions of nanodomain radii for SRRM2 inside (orange) and outside (blue) nuclear speckles (bars). Solid lines show two-Gaussian fits to the distributions. Inset, mean nanodomain radii for SRRM2 inside and outside speckles. n = 4070 nanodomains for SRRM2 inside and 4670 nanodomains for SRRM2 outside. **d,** Grouping of consecutive trajectory segments into confined and diffusive periods. Top, schematic showing individual confined periods in distinct colors and diffusive periods in black. Bottom, representative SRRM2 trajectory inside a nuclear speckle, with individual confined periods shown in distinct colors and diffusive periods shown in black. **e,** Distributions of diffusive-state dwell times for SRRM2 inside (orange) and outside (blue) nuclear speckles. Diamond markers show the observed distributions, and solid lines show fits to a single-exponential function for dwell times >5 ms. Inset represents mean diffusive-state dwell times inside and outside speckles. n = 3495 diffusive periods for SRRM2 inside and 7555 diffusive periods for SRRM2 outside. **f,** Distributions of confined-state dwell times for SRRM2 inside (orange) and outside (blue) nuclear speckles. Diamond markers show the observed distributions, and solid lines show fits to a Lomax distribution. Inset represents mean confined-state dwell times inside and outside speckles. n = 3639 confined periods for SRRM2 inside and 7748 confined periods for SRRM2 outside. **g,** Probability density functions of the confined-state exit rate k for SRRM2 inside (orange) and outside (blue) nuclear speckles, obtained from the Lomax fits. **h,** The mean diffusive displacement between adjacent nanodomains of SRRM2 (red), SON (purple) and SRSF1 (blue) inside (solid) and outside (diagonal hatching) nuclear speckles. n = 3495, 7555, 9060, 5835, 3092, 4777 diffusive periods, from left to right. Bar graphs in (**c**, **e**–**f**, **h**) show mean ± SEM. (*) p ≤ 0.05, (**) p ≤ 0.01, and (***) p ≤ 0.001 (Mann-Whitney U test). **i,** Higher-order spatial organization of nanodomains within a SRRM2 trajectory inside a nuclear speckle. Nanodomains were clustered using the DBSCAN algorithm, with colors indicating distinct higher-order assemblies. **j**, The mean assembly area of SRRM2 (red), SON (purple), and SRSF1 (blue) inside nuclear speckles. n = 343, 586, 306 assemblies, from left to right. Data are presented as mean ± SEM. (*) p ≤ 0.05, (**) p ≤ 0.01, and (***) p ≤ 0.001 (Mann–Whitney U test). **k,** Distribution of the number of nanodomains per assembly for SRRM2 inside nuclear speckles (bars), fitted to a logarithmic distribution (solid line). n = 343 assemblies.

For a dwell-time analysis of the multi-state diffusive behavior, we grouped consecutive confined or diffusive segments together (Fig.4d). Dwell times of diffusive periods primarily followed an exponential distribution across all three proteins (mean ∼7 ms, with rare events extending up to ∼50 ms for SRRM2 inside speckles), indicating a kinetically homogeneous exit from the diffusive state (Fig.4e and Extended Data Fig.4d–e). In contrast, dwell times of confined periods, although dominated by short events, exhibited a long tail across all three proteins (mean ∼50 ms for SRRM2 inside speckle), indicating heterogeneous confinement dynamics (Fig.4f and Extended Data Fig.4f–g). Consistent with this heterogeneity, the rate *k* of leaving the confined state also displayed a broad distribution, indicating that molecules within the confined state exhibit a range of exit kinetics (Fig.4g and Extended Data Fig.4h–i). Most confinement periods were short-lived, consistent with transient interactions, whereas a small fraction persisted much longer, suggesting more stable associations with the local molecular environment (Fig.4f–g and Extended Data Fig.4f–i). To characterize molecular movement between successive confinement periods, we measured the displacement during each diffusive segment before entry into the next confined period. On average, molecules traveled ∼80 nm between successive confined periods within the speckle for all three proteins (Fig.4h). These results suggest that speckle proteins repeatedly alternate between a kinetically homogeneous diffusive state and a kinetically heterogeneous confined state. Whereas diffusion between speckle nanodomains follows relatively uniform kinetics, the broad distribution of confinement lifetimes is consistent with a spectrum of local interactions, potentially ranging from weak, transient encounters to more stable multivalent associations.^74^

Surprisingly, although all three speckle proteins were more diffusive outside nuclear speckles, their trajectories still contained transient confined states (Fig.4c and Extended Data Fig.4a–b). This suggests that confinement is not unique to the high-density speckle environment but can also arise from transient interactions occurring outside the speckles, even at lower molecular concentrations. Yet, the nanodomains observed inside and outside speckles exhibited distinct physical properties. First, molecules explored larger confined regions outside the speckle, with the average confinement radius being larger for all three proteins (Fig.4c and Extended Data Fig.4a–b). Second, the scaffold proteins SON and SRRM2 exhibited longer diffusive dwell times and shorter confined dwell times outside speckles, indicating weaker and more transient associations with the local molecular environment (Fig.4e–f and Extended Data Fig.4d,f). SRSF1 showed shorter confined dwell times but similar diffusive dwell times outside speckles (Extended Data Fig.4e,g). Third, the nanodomains were more sparsely distributed outside speckles, and molecules traveled longer distances between successive confined periods (Fig.4h). Together, these physical properties imply that confinement outside speckles is governed by a sparser and less interconnected interaction landscape, in which reduced local interaction density gives rise to larger confinement regions, weaker transient associations, and increased spacing between speckle nanodomains.

Within individual trajectories, proteins transitioned among neighboring speckle nanodomains before making occasional long-range excursions (Fig.4a). This behavior suggested that nanodomains are not randomly distributed throughout nuclear speckles but are instead spatially organized into locally enriched assemblies. To test this possibility, we grouped neighboring speckle nanodomains using DBSCAN algorithm to identify higher order nanodomain assemblies (Fig.4i, see Methods). These higher-order assemblies define larger spatial regions within which the molecules repeatedly transition among neighboring speckle nanodomains before undergoing long-range diffusion. Spatially, the observed SRRM2 nanodomain assemblies cover ∼ 0.02μm^2^ (∼ 0.015μm^2^ for SRSF1 and ∼ 0.03μm^2^ for SON) inside the speckles (Fig.4j, see Methods). Within the assemblies the distances between centroids of neighboring speckle nanodomains were ∼70 nm inside speckles for all three proteins, and were significantly larger outside speckles, consistent with a more sparsely populated environment (Extended Data Fig.4j). We hypothesize that the higher-order spatial organization of speckle nanodomains likely emerges from multivalent intermolecular interactions that generate regions of elevated interaction frequency and stability, allowing molecules to hop between nearby speckle nanodomains before escaping to more distant locations.

To determine whether speckle nanodomain assemblies exhibit a similar hierarchical organization to the molecular clusters identified in fixed-cell MINFLUX super-resolution imaging, we quantified the number of observed speckle nanodomains within each assembly. For all three proteins, the resulting assembly-size distributions were well described by a logarithmic distribution, consistent with the heterogeneous cluster organization observed in the MINFLUX super-resolution imaging of fixed cells (Fig.4k and Extended Data Fig.4k–m). Within nuclear speckles, we observed that each assembly contains on average ∼3.6 speckle nanodomains for SRRM2, ∼3.3 for SRSF1, and ∼5 for SON (Fig.4k and Extended Data Fig.4l–m). While most of the assemblies were small, consistent with the transient nature of local interactions, a subset of assemblies expanded into larger assemblies, resulting in a heavy-tailed cluster-size distribution. Such heavy-tailed distribution suggests that higher-order organization emerges through stochastic association of neighboring nanodomains rather than assembly into structures with fixed dimensions, generating hierarchical spatial organization.

Together, our results converge on a model in which nuclear speckles exhibit a hierarchical, two-layered organization composed of nanoscale interaction domains, referred to as speckle nanodomains, embedded within larger dynamic assemblies. Individual proteins repeatedly transition between neighboring speckle nanodomains before escaping to distant clusters, directly linking the heterogeneous architecture of speckles to their molecular dynamics. Such multiscale organization resembles the hierarchical structure of microphase-separated splicing factor condensates that we observed *in vitro*,^13^ suggesting that similar physical principles may organize nuclear speckles *in vivo*.

### Condensate remodeling reshapes nanoscale interaction environments

To determine whether this hierarchical architecture responds dynamically to functional perturbations, we evaluated speckle protein and RNA organization and dynamics following treatment with the transcription inhibitor DRB, splicing inhibitor PlaB, L-arginine to perturb charge-mediated interactions, depletion of SON, SRRM2 or *MALAT1*, or acute hypoxia (0.2% O₂ or cobalt chloride for 24 h) (Fig.5, Extended Data Fig.5, Supplementary Tables 1–5).

**Fig. 5:**
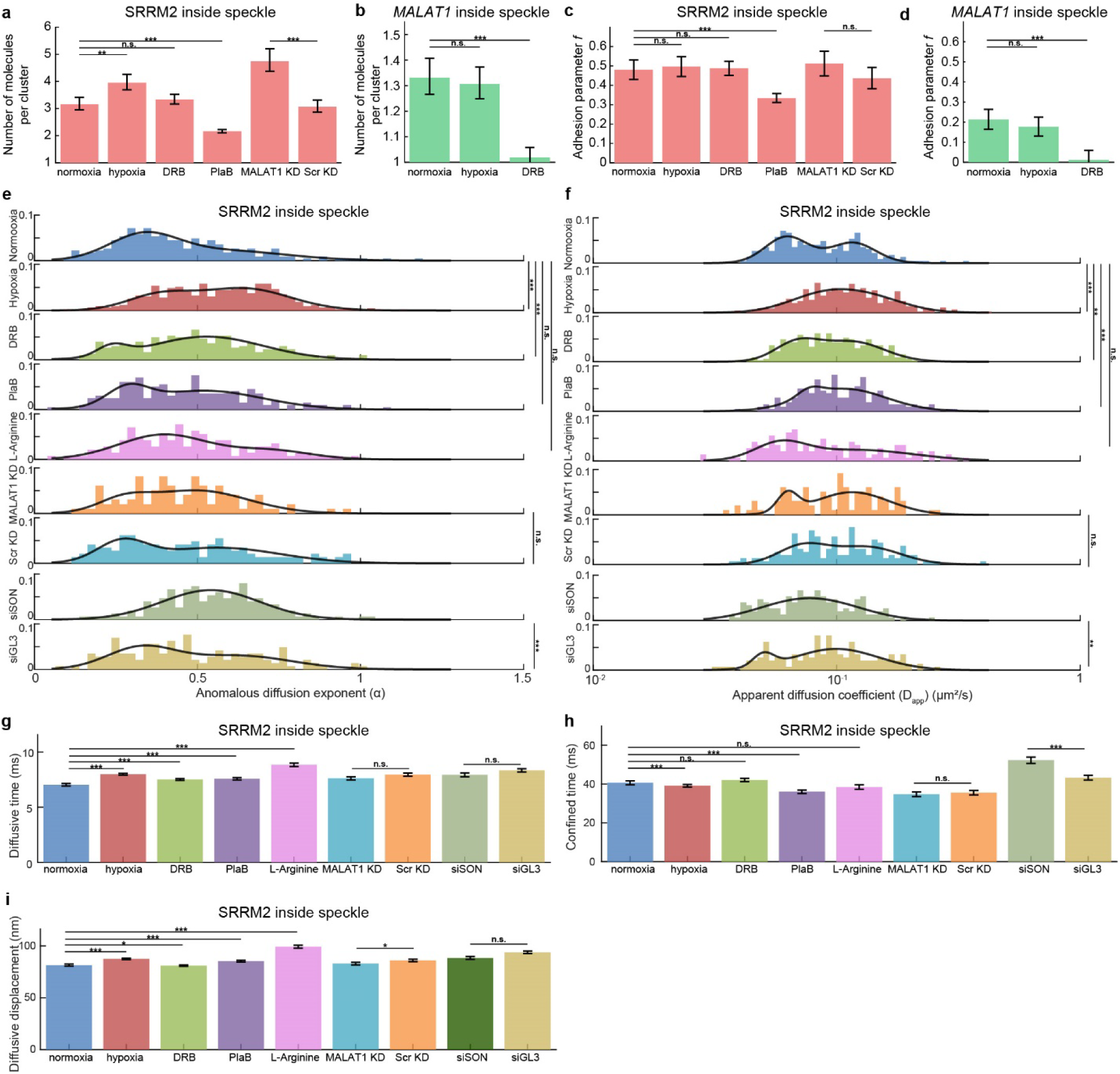
Perturbations alter the spatial organization and dynamics of speckle components. **a–b,** The expected number of molecules per cluster for SRRM2 (**a**) and *MALAT1* (**b**) inside nuclear speckles after different perturbations. **c–d,** Adhesion parameter *f* for SRRM2 (**c**) and *MALAT1* (**d**) inside nuclear speckles after different perturbations. Bar graphs in (**a-d**) show means with 95% confidence intervals (CIs). Asterisks denote non-overlapping CIs: (*) 95%, (**) 99%, (***) 99.9%; n.s. denotes overlapping 95% CIs. **e,** Distributions of the anomalous diffusion exponent (α) for SRRM2 inside nuclear speckles after different perturbations. **f,** Distributions of the apparent diffusion coefficient (D_app_) for SRRM2 inside nuclear speckles under perturbations. The distributions in (**e**) and (**f**) are fitted to a sum of two Gaussians as a guide to the eye (solid lines). (*) p ≤ 0.05, (**) p ≤ 0.01, and (***) p ≤ 0.001 (Mann-Whitney U test). α and D_app_ were obtained from the same trajectories. n = 225, 431, 304, 199, 143, 98, 146, 150, and 144 trajectories, from top to bottom. **g–h,** The mean diffusive-state dwell times (**g**) or confined-state dwell times (**h**) of SRRM2 after different perturbations. **i,** The mean diffusive displacement of SRRM2 after different perturbations. Bar graphs in (**g**–**i**) show mean ± SEM. (*) p ≤ 0.05, (**) p ≤ 0.01, and (***) p ≤ 0.001 (Mann– Whitney U test). For (**g**) and (**i**), n = 3495, 9425, 5847, 4164, 2481, 2233, 2554, 1763, and 2546 diffusive periods, from left to right. For (**h**), n = 3639, 9578, 5992, 4356, 2644, 2348, 2696, 1920, and 2690 confined periods, from left to right.

Across all perturbations, both MINFLUX fixed cell imaging as well as live-cell tracking revealed remodeling of molecular clustering, nanodomain organization, and diffusion dynamics (Fig.5 and Extended Data Fig.5), indicating that higher-order interaction environments are highly sensitive to perturbation-induced changes in speckle nanodomain organization. A comprehensive summary of the effects of each perturbation on these organizational features for all proteins and RNAs analyzed is provided in Supplementary Tables 1–5. Notably, different perturbations did not simply increase or decrease molecular abundance but selectively remodeled distinct organizational features of the speckle nanodomains. Changes in organizational features and dynamics frequently occurred independently of protein and RNA concentration, demonstrating that nanodomain composition alone does not determine its organization or molecular dynamics (Supplementary Tables 1–5).

For instance, acute hypoxia and splicing inhibition illustrate two fundamentally distinct modes of condensate remodeling. Splicing inhibition broadly destabilized hierarchical organization of nuclear speckles by reducing SRRM2 concentration, cluster size and level of molecular adhesion, indicating impaired nanoscale scaffold assembly (Fig.5a,c and Extended Data Fig.5c,f). At the same time, SRRM2 exhibited increased molecular mobility, reflected by a higher D_app_, together with prolonged diffusive dwell times, increased diffusive distances and shortened confined dwell times, indicating enhanced molecular exchange between speckle nanodomains and alteration of the interaction network that maintains speckle architecture (Fig.5e–i). In contrast, hypoxia promoted the assembly of larger SRRM2 clusters while simultaneously increasing SRRM2 molecular mobility, with increased D_app_ and α, and destabilizing speckle nanodomain assembly through altered dwell times and increased diffusive distance (Fig.5a,c,e–i and Extended Data Fig.5a,f). These contrasting responses demonstrate that enhanced molecular enrichment can coexist with a more dynamic interaction environment and that condensate state should not be inferred solely based on molecular abundance.

Additional perturbations further demonstrated that different regulatory pathways remodel distinct organizational features of the speckle nanodomain assembly. For instance, transcription inhibition by DRB treatment increased SRRM2 mobility and destabilized speckle nanodomains without substantially altering higher-order condensate organization (Fig.5a,c,e–h and Extended Data Fig.5b,f). In contrast, depletion of SON increased the α value of SRRM2 while decreasing its D_app_ and prolonging confined dwell times, indicating that the degree of molecular constraint and overall mobility can change independently (Fig.5e–h). L-arginine treatment, which was previously shown to weaken microphase separation of splicing factors *in vitro*,^13^ increased the diffusive dwell times and diffusive distances without altering overall SRRM2 mobility, consistent with a sparser spatial distribution of speckle nanodomains (Fig.5e–i). Conversely, *MALAT1* depletion promoted formation of larger SRRM2 assemblies without changing mobility (Fig.5a,c,e–h and Extended Data Fig.5d–f). Together, these findings reveal that different molecular perturbations remodel specific speckle-resident proteins differently rather than producing a common structural response. This could be indicative of heterogeneities in the interaction networks that generate distinct speckle nanodomains.

Similarly, speckle-associated RNAs responded with species-specific selectivity: DRB caused *MALAT1* to redistribute into the nucleoplasm, reducing its concentration, molecular adhesion *f*, and cluster size inside speckles (Fig.5b,d and Extended Data Fig.5b,f). *MALAT1* depletion reduced both the *U2 snRNA* concentration and cluster size, whereas hypoxia decreased *U2 snRNA* cluster size without altering its concentration inside speckles (Extended Data Fig.5f–h). These observations show that nanodomain properties, higher-order nanodomain assembly, and molecular dynamics could be selectively remodeled by physiological, pharmacological, and genetic perturbations, rather than changing through a uniform condensate-wide response.

## Discussion

Biomolecular condensates have emerged as a fundamental principle of cellular organization. *In vitro* studies have revealed nanoscale and hierarchical organization within biomolecular condensates.^5, 10, 12, 13, 72, 75^ For instance, using single-molecule tracking in reconstituted FUS condensates, Walter and colleagues recently identified nanometer-scale domains that locally confine RNA and protein diffusion and regulate molecular residence times, supporting a heterogeneous, percolated-network model of condensate interiors.^10, 72^ However, direct evidence for similar organization in living cells, and its impact on molecular dynamics, has remained lacking. Using MINFLUX nanoscopy for both quantitative single-molecule imaging and live-cell tracking, we provide direct *in vivo* evidence that nuclear speckles exhibit dynamic, heterogeneous organizations rather than behaving as homogeneous liquid compartments. Speckle components formed nanoscale molecular assemblies, with individual protein molecules exhibiting heterogeneous mobility and transient confinement within ∼40 nm radius nanodomains, and the neighboring nanodomains were further organized into higher order assemblies. Our findings establish a quantitative framework, connecting nanoscale molecular organization, transient confinement, and dynamic exchange within a living biomolecular condensate.

Our quantitative molecular inventory provides estimates of the absolute concentrations of several nuclear speckle components inside and outside nuclear speckles (Fig.1e). For SRRM2, our estimate is broadly consistent with a fluorescence-based measurement of endogenous SRRM2 concentration, despite differences in cell type and quantification approach.^18^ Previous studies have also reported cellular abundance or copy-number estimates for several other speckle components, primarily using quantitative mass spectrometry.^61^ In contrast to whole-cell abundance measurements, our measurements provide estimates of local concentrations inside and outside nuclear speckles. Direct quantitative comparison between these approaches is complicated by differences in their underlying measurements and assumptions. Our absolute concentration estimates rely on converting two-dimensional MINFLUX measurements to volumetric concentrations using an assumed effective axial sampling depth (see Methods), which introduces uncertainty into the resulting concentration estimates. Despite these uncertainties, the measurements provide a quantitative concentration framework for interpreting the nanoscale spatial organization of speckle components.

Our study further extends models of nuclear speckle assembly beyond mesoscale compartmentalization.^18, 42^ Recent studies demonstrated that the scaffold proteins SRRM2 and SON form immiscible sub-compartments within nuclear speckles, with SRRM2 phase separation and higher-order oligomerization driving sub-compartment assembly and SON occupying a distinct scaffold phase.^18, 19, 76^ At the molecular scale, our analyses revealed non-random local association of speckle components. By extending the hard-disk model to include an effective adhesion parameter, we quantified this tendency for each component. The inferred adhesion parameter provides a phenomenological measure of local molecular adhesion and likely reflects, at least in part, homotypic interactions among molecules of the same species. Notably, SRRM2 and SON exhibited the highest inferred adhesion parameters among the speckle components examined (Fig.2g), consistent with their proposed scaffold roles. Our results therefore extend the previous observations by quantitatively describing molecular organization below the scale of optical sub-compartments.

Molecular mobility within biomolecular condensates is often described as diffusion through a relatively continuous and homogeneous environment.^1–3, 12^ In contrast, our live-cell tracking data revealed substantial heterogeneity in the motion of individual speckle proteins at sub-millisecond timescales, as reflected by the broad distributions of α and D. The generally lower mobility within speckles further indicates that the local speckle environment constrains molecular motion. Resolving dynamics at sub-millisecond timescales provides insight into how molecules explore their immediate surroundings and may capture transient intermolecular interactions that contribute to the dynamic molecular assemblies within nuclear speckles. Analysis of individual trajectories further revealed that this heterogeneous motion consisted of alternating diffusive and confined periods. During confined periods, molecules explored nanodomains of ∼40 nm in radius, with broad dwell-time distributions revealing substantial heterogeneity in the persistence of molecular confinement. Moreover, individual molecules transitioned among neighboring nanodomains, which were organized into higher-order assemblies. Nuclear speckles have been proposed to locally enrich splicing factors before their recruitment to actively transcribed genes.^69^ When confined within nanodomains, molecules explore a restricted spatial range, which may increase the probability of local molecular interactions. Meanwhile, molecules can transition among neighboring nanodomains and undergo intervening diffusive periods, allowing them to sample different local environments. Such dynamic organization may therefore promote local homotypic and heterotypic molecular encounters while maintaining the molecular exchange required for speckle function.

Static MINFLUX imaging and live-cell tracking provide complementary views of nanoscale speckle organization, revealing spatial molecular organization and molecular dynamics, respectively. The comparable length scales of molecular clusters detected by static imaging and nanodomains identified from confined molecular motion in live cells suggest a relationship between local molecular association and transient confinement, although these measurements describe distinct properties and should not be considered geometrically equivalent. Notably, inferred adhesion parameters differed substantially among speckle proteins, whereas nanodomain dimensions and confinement dynamics were comparatively similar across species. The adhesion parameter describes effective spatial associations only among molecules of the same species, suggesting that homotypic interactions alone are unlikely to fully account for the observed nanodomain organization and dynamics. Instead, these properties likely emerge from a broader network of homotypic and heterotypic protein-protein and protein-RNA interactions. Such a network-based organization is consistent with emerging models proposing that biomolecular condensates behave as heterogeneous and percolated interaction networks rather than homogeneous equilibrium liquid phases.^5, 9, 10, 12^ The ability to visualize these interaction environments in living cells provides an experimental framework for linking molecular interactions to condensate material properties across multiple spatial scales.

Our findings bridge *in vitro* observations of microphase separation with *in vivo* nuclear architecture. Splicing factors and *MALAT1* reconstituted *in vitro* form stable nanoscale microphases that subsequently organize into larger mesoscale clusters.^13^ The speckle nanodomains identified here resemble these microphases in spatial dimension, higher-order assembly behavior, and sensitivity to electrostatic disruption by L-arginine. Together with our previous *in vitro* findings that *MALAT1* drives SRSF1 microphase separation,^13^ these results support the possibility that related interaction principles contribute to nanoscale organization in both reconstituted SRSF1-*MALAT1* assemblies and in nuclear speckles. The *MALAT1*-depletion phenotypes further indicate that *MALAT1* modulates the organization of selected speckle components, although it is not required for all speckle features, including nanodomain assembly or splicing factor dynamics.^20^

Finally, our findings suggest that condensate organization is encoded not only by molecular composition but also by the interaction networks underlying its hierarchical organization. Across perturbations, changes in molecular adhesion, nanodomain organization, confinement, and molecular mobility were not consistently coupled to changes in protein concentration or to one another. These observations suggest that condensate composition, spatial organization, and molecular dynamics represent distinct but interconnected dimensions of condensate regulation.

An important strength of this study is the ability to resolve nanoscale organization and molecular dynamics within living biomolecular condensates, while also identifying questions for future investigation. Although we identify dynamic speckle nanodomains as the organizational units of nuclear speckles, their molecular composition and stoichiometry remain unknown. Determining whether different nanodomains contain distinct combinations of proteins and RNAs will require multiplexed imaging at comparable spatial resolution. The molecular mechanisms by which specific protein-protein and protein-RNA interactions generate, stabilize, and remodel speckle nanodomains remain to be established. Finally, our work provides a physical framework for understanding how condensate architecture may regulate RNA metabolism, but it does not directly link individual nanodomains to transcriptional or splicing outputs. Integrating nanoscale imaging with real-time measurements of gene expression and targeted perturbations of condensate architecture will be necessary to establish these causal relationships. Currently, such key questions have remained experimentally inaccessible because the relevant spatial and temporal scales could not be resolved in living cells. By providing direct visualization of condensate architecture and molecular dynamics at nanometer and millisecond resolution, our work establishes an experimental framework for addressing these longstanding mechanistic questions.

## Resource availability

### Correspondence

Further information and requests for resources and reagents should be directed to the lead contacts, Kannanganattu V. Prasanth, Ido Golding; Taekjip.Ha

### Materials availability

Unique materials generated by this study, including cell lines, are available upon request from Kannanganattu V. Prasanth.

### Data and code availability

Any additional information required to reanalyze the data reported in this manuscript is available from the lead contacts upon request.

## Acknowledgements

We thank Profs. Rohit Pappu (Wash. U.) and Ashish Lal (NCI) for their insightful comments. We thank Dr. Anders S Hansen (MIT) for reagents and help. We thank Jessica Matthias, Otto Wirth, Kingsley Boateng, Reza Rajabi and Glenn Fried for technical assistance with the MINFLUX microscope in IGB. We thank Joseph Slivka and Sepehr Alaeen for suggestions on MINFLUX tracking.

## Funding

This work was supported by grants from NSF Science and Technology Center for Quantitative Cell Biology (NSF DBI 2243257) to KVP, SGP, IG, & TH, SGP (NSF 2225464); NIH R01GM132458 to KVP, 1R35GM152450 to SGP, R01 EY012141 to SHD, R35GM140709, and the Alfred P. Sloan Foundation grant (G-2023-19649) to IG, and ARPA-H AY1AX000030 to KVP. J.Z. was supported by a Feodor Lynen Research Fellowship from the Alexander von Humboldt Foundation.

## Author contributions

ML, TY, JZ, TH, IG and KVP conceived and led the project. ML designed and performed all MINFLUX imaging and tracking, TY analyzed the MINFLUX imaging data, JZ analyzed the MINFLUX tracking data and performed wide-field SPT. YJS, QM, and ADH designed and generated the KI cells. SHD and DF helped in image data acquisition and analyses. SMF designed and generated the control and *MALAT1* ASOs. ML, TY, JZ interpreted the data with help from SGP, IG, TH and KVP. ML, TY and JZ composed the manuscript with editorial input from IG, TH and KVP. All authors read and commented on the manuscript.

## Competing interests

SMF is an employee of Ionis Inc. and receives a salary.

## Declaration of AI-assisted writing

AI tools were used in portions of the manuscript to improve grammar and conciseness. After using this tool, the authors reviewed and edited the content as needed and take full responsibility for the content of the published article.

## Extended Data figures

**Extended Data Fig. 1.**
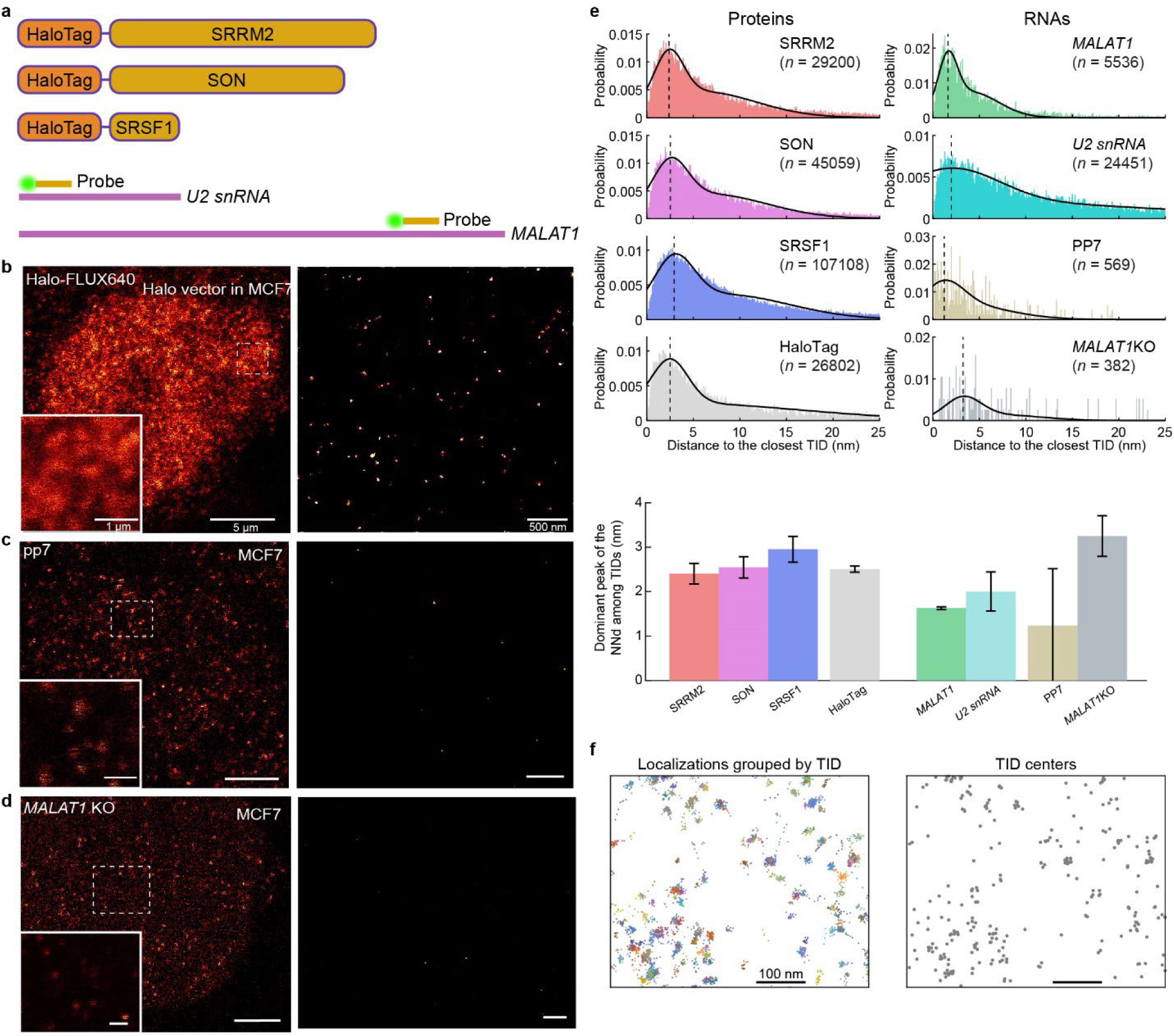
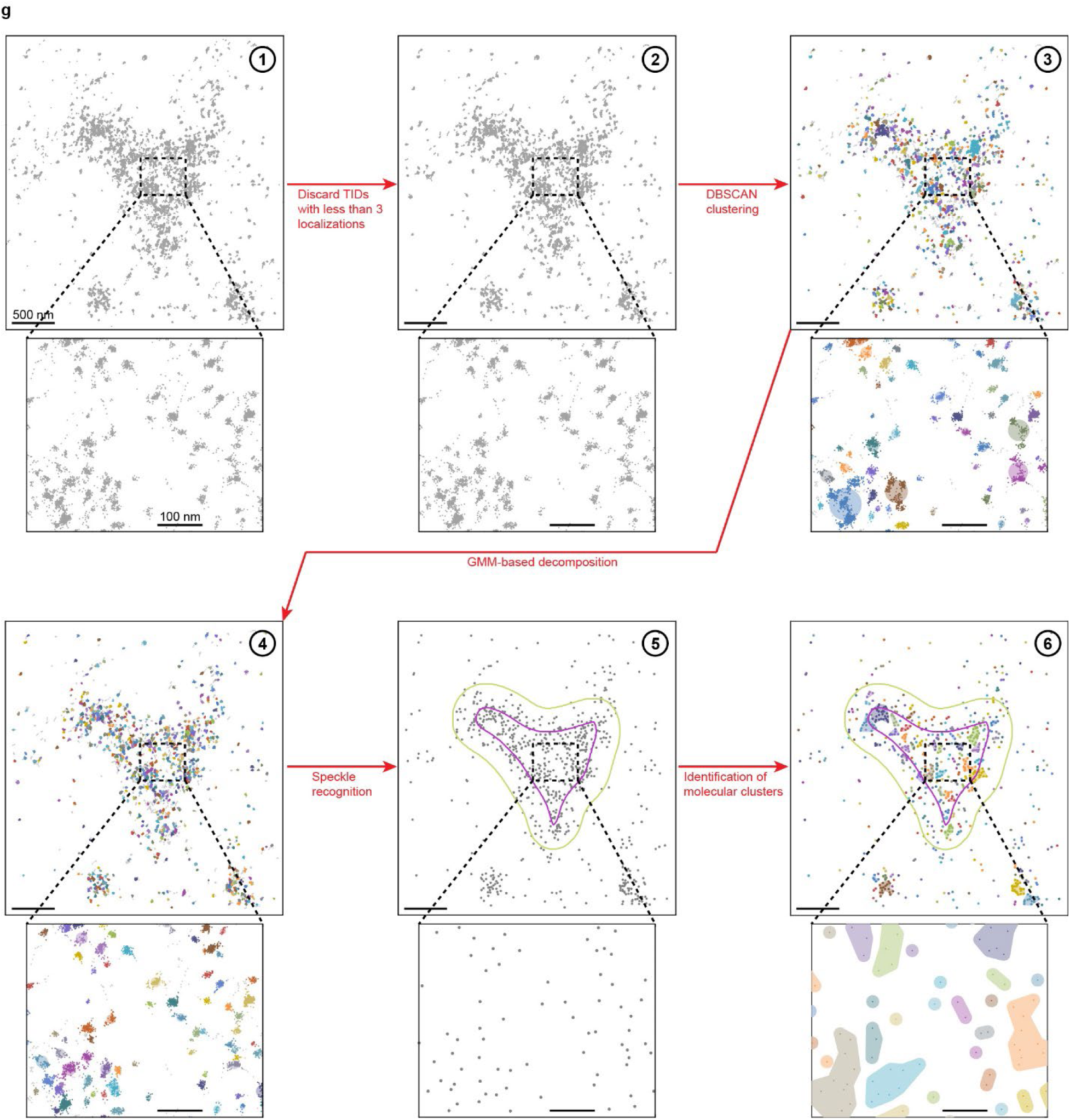
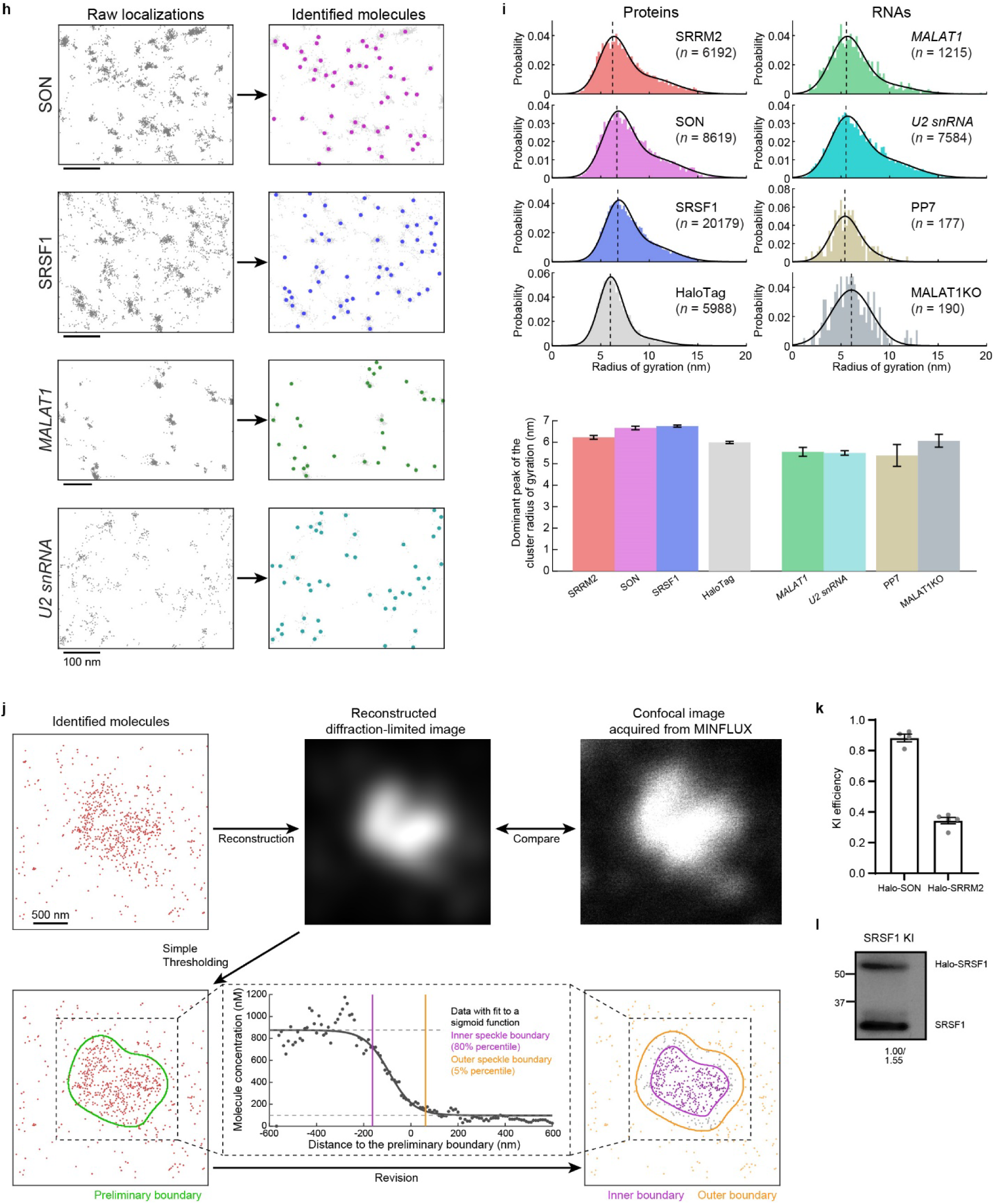
Labeling strategies and analysis workflow for MINFLUX imaging. **(a)** Labeling strategies for speckle proteins and RNAs. Top, endogenous speckle protein genes are N-terminally fused to HaloTag (with a 9 amino acid linker in between) and labeled with FLUX-640 HaloTag ligand. Bottom, speckle-resident *U2 snRNA* and *MALAT1* RNAs are labeled by smFISH using ∼45-nt oligonucleotide probes conjugated to sCy5 NHS ester, with a single probe hybridized to each RNA molecule. Schematics are not drawn to scale. **(b–d)** Representative 2D MINFLUX images of control samples. (**b**) Exogenously expressed HaloTag labeled with FLUX-640 HaloTag ligand, serving as a control for HaloTag-based protein labeling. (**c**) WT MCF7 cells labeled with PP7-targeting smFISH probes. (**d**) *MALAT1*-knockout MCF7 cells labeled with a *MALAT1*-targeting smFISH probe. (**c**) and (**d**) serve as controls for smFISH-based RNA labeling. Left, confocal images acquired on the MINFLUX microscope showing the nucleus. Inset, magnified view of the selected region used for MINFLUX imaging. Right, 2D MINFLUX images of the corresponding regions. **(e)** Top, nearest-neighbor distance distributions between MINFLUX trace positions for SRRM2, SON, SRSF1, *MALAT1*, *U2 snRNA*, and the control samples (HaloTag alone, PP7-targeting probes, and *MALAT1*-knockout cells). Each trace position is defined as the geometric center of the localizations sharing the same TID. Colored bars show the observed distributions, and solid lines show fits to a sum of two Gaussians. Dashed lines indicate the fitted peak position of the dominant Gaussian component. Bottom, summary of the peak positions of the dominant Gaussian components for all samples. Bars show the fitted peak positions, with error bars indicating 95% confidence intervals from the two-Gaussian fits. **(f)** Representative visualization of TIDs in an SRRM2 MINFLUX dataset. Left, MINFLUX localizations grouped based on TID, shown in distinct colors. Right, corresponding trace positions, shown as gray dots. **(g)** MINFLUX image analysis workflow, illustrated using a representative SRRM2 dataset. (1) Raw MINFLUX localizations. (2) Localizations retained after discarding TIDs containing fewer than three localizations. (3) DBSCAN grouping of the retained localizations, with colors indicating individual DBSCAN clusters and circles centered on each cluster with radii corresponding to their radii of gyration. (4) Result after GMM-based decomposition of selected DBSCAN clusters, shown for the complete dataset. Colors indicate the resulting clusters, and circles are centered on each cluster with radii corresponding to their radii of gyration. The resulting clusters were interpreted as individual molecules. (5) Determination of nuclear speckle boundaries from the identified molecular positions, with the inner and outer boundaries shown in purple and yellow, respectively. (6) Molecular clusters identified by a second round of DBSCAN. Individual molecular clusters are shown in distinct colors, with corresponding shaded regions added for visualization. **(h)** Identification of individual SRRM2, SRSF1, *MALAT1*, and *U2 snRNA* molecules from MINFLUX localizations. Left, raw MINFLUX localizations. Right, identified molecule positions (colored) overlaid on the corresponding MINFLUX localizations (gray). **(i)** Top, distributions of cluster radii of gyration for SRRM2, SON, SRSF1, *MALAT1*, *U2 snRNA*, and the control samples (HaloTag alone, PP7-targeting probes, and *MALAT1*-knockout cells). Colored bars show the observed distributions, and solid lines show fits to a sum of two Gaussians. Dashed lines indicate the fitted peak position of the dominant Gaussian component. Bottom, summary of the peak positions of the dominant Gaussian components for all samples. Bars show the fitted peak positions, with error bars indicating 95% confidence intervals from the two-Gaussian fits. **(j)** Identification of nuclear speckle boundaries based on local molecular density, illustrated using a representative SRRM2 dataset. A diffraction-limited image is reconstructed by rendering the identified molecule positions as two-dimensional Gaussian functions and compared with the corresponding confocal image acquired on the MINFLUX microscope. A preliminary speckle boundary is obtained by thresholding the reconstructed image. Local molecular concentration is then quantified directly from the identified molecular positions as a function of signed distance from the preliminary boundary and fitted with a sigmoid function. The fitted concentration profile is used to determine the inner and outer speckle boundaries. **(k)** Knock-In efficiencies of the Halo-SON and Halo-SRRM2 cell lines quantified by ddPCR. Individual points represent independent measurements of biological replicates, and bars show the mean ± SEM. **(l)** Knock-In efficiency of the Halo-SRSF1 cell line quantified by Western blotting using SRSF1 antibody. Halo-SRSF1 and endogenous SRSF1 are resolved as the upper and lower bands, respectively. Numbers below the blot indicate relative band intensities, which were used to estimate the knock-in efficiency.

**Extended Data Fig. 2.**
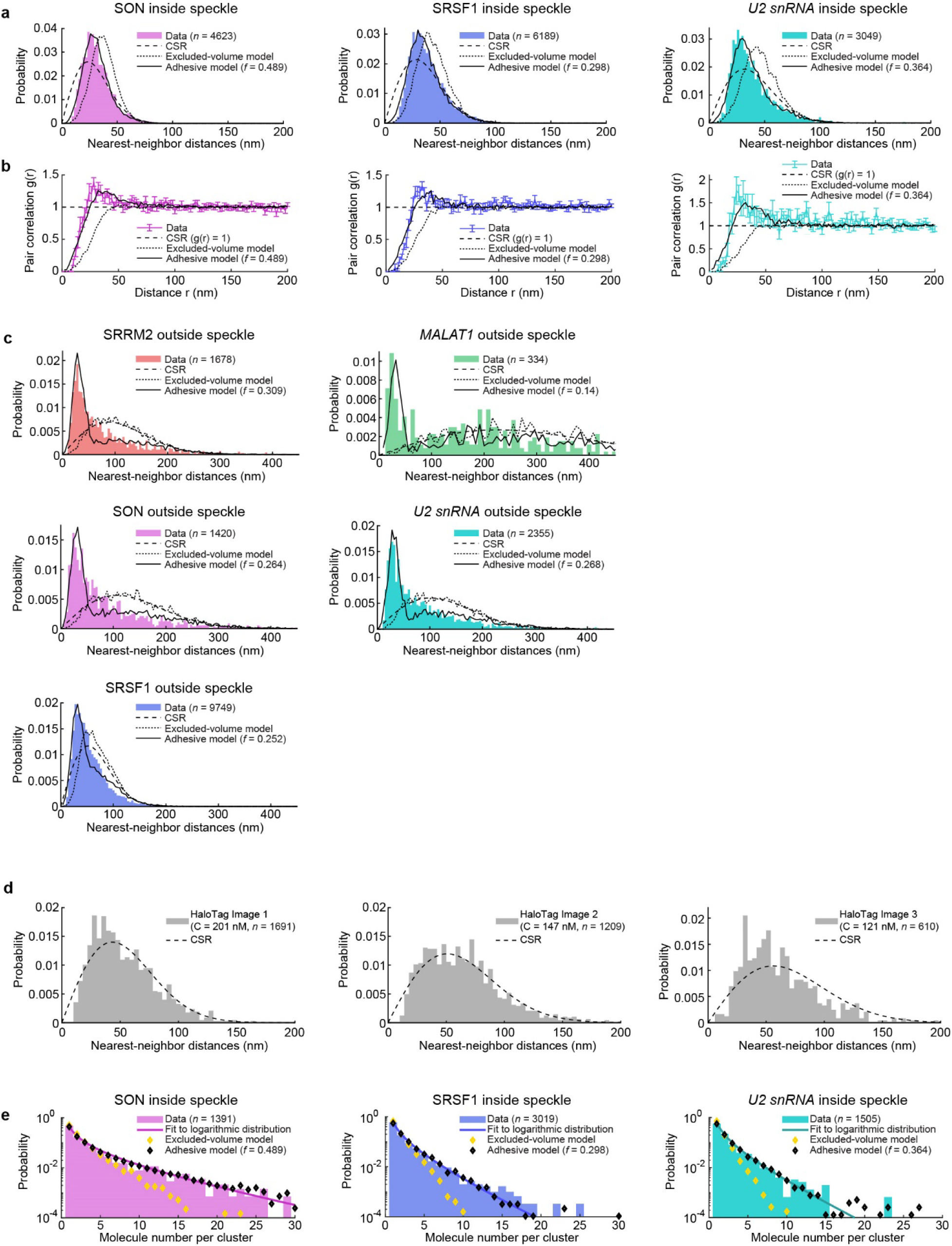
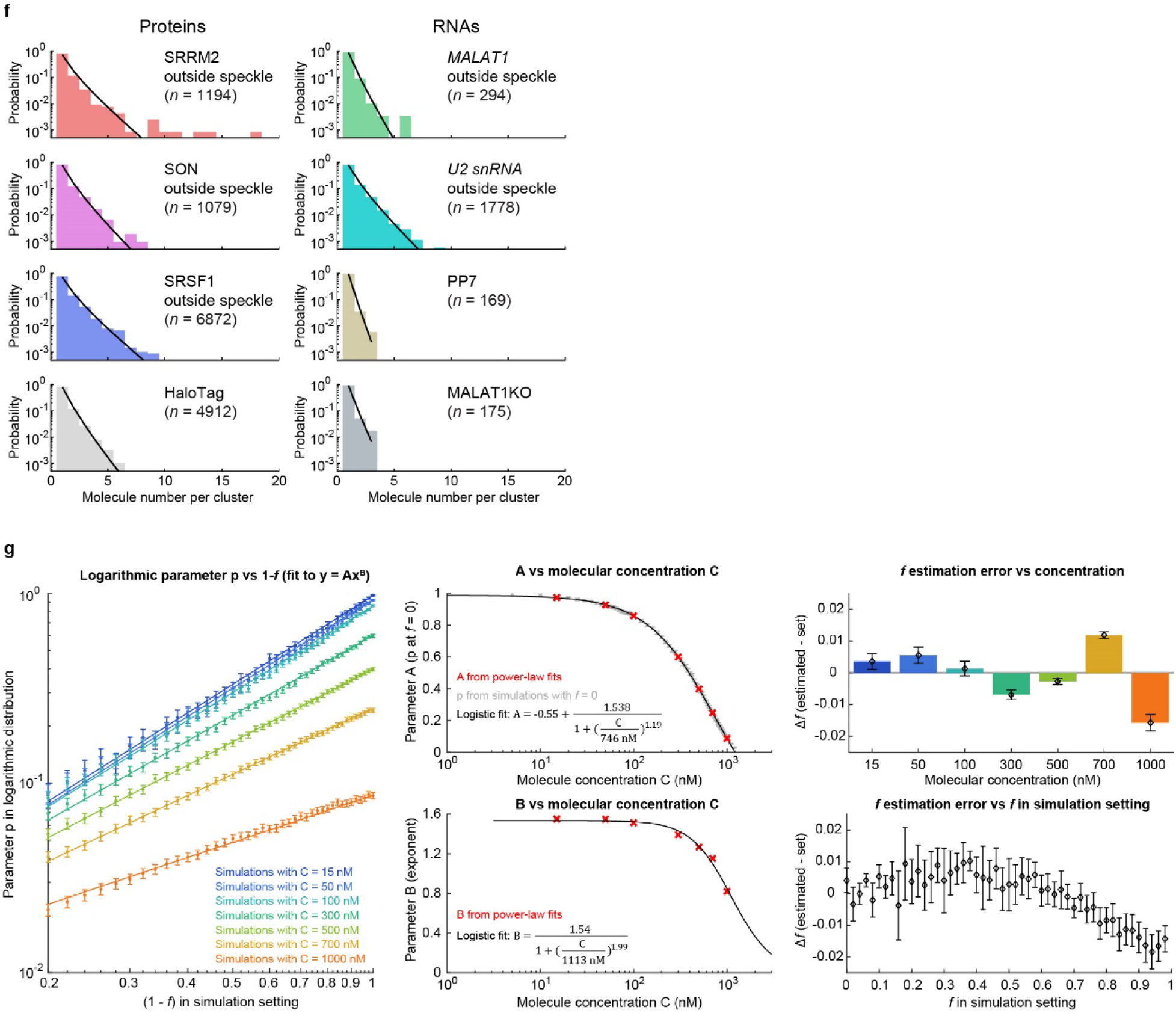
Quantitative analysis and simulation of molecular clustering. **(a)** Nearest-neighbor distance distributions of SON (left), SRSF1 (middle), and *U2 snRNA* (right) molecules inside nuclear speckles, pooled across speckles (bars), compared with complete spatial randomness (CSR; dashed lines), the excluded-volume model without local molecular adhesion (dotted lines), and the excluded-volume model with local molecular adhesion (solid lines). **(b)** Pair correlation functions, g(r), of SON (left), SRSF1 (middle), and *U2 snRNA* (right) molecules inside nuclear speckles, averaged across speckles (colored lines with error bars), compared with CSR (dashed lines), the excluded-volume model without local molecular adhesion (dotted lines), and the excluded-volume model with local molecular adhesion (solid black lines). **(c)** Nearest-neighbor distance distributions of SRRM2, SON, SRSF1, *MALAT1*, *U2 snRNA* localized outside nuclear speckles, pooled across images (bars), compared with CSR (dashed lines), the excluded-volume model without local molecular adhesion (dotted lines), and the excluded-volume model with local molecular adhesion (solid lines). **(d)** Nearest-neighbor distance distributions of HaloTag-alone molecules computed separately for individual images (bars), each compared with CSR simulated at the corresponding molecular concentration (dashed lines). **(e)** Distributions of the number of molecules per cluster for SON (left), SRSF1 (middle), and *U2 snRNA* (right) inside nuclear speckles, pooled across speckles (bars) and fitted to a logarithmic distribution (solid lines), compared with simulations from the excluded-volume model without local molecular adhesion (yellow markers) and the excluded-volume model with local molecular adhesion (black markers). **(f)** Distributions of the number of molecules per cluster for SRRM2, SON, SRSF1, *MALAT1*, *U2 snRNA* outside nuclear speckles and the three control samples (HaloTag alone, PP7-targeting probes, and *MALAT1*-knockout cells) (colored bars). Each dataset is fitted to a logarithmic distribution (solid lines). **(g)** Empirical mapping between the adhesion parameter *f* and the logarithmic fit parameter *p*. For each simulated dataset spanning a range of *f* and molecular concentrations C, the cluster-size distribution is fitted with a logarithmic distribution to obtain *p*. Left, relationship between *p* and 1-*f* at seven simulated molecular concentrations, fitted with *p* = A(1-*f*)^B^. Middle, dependence of A (top) and B (bottom) on molecular concentration C. Red crosses show A and B obtained from the power-law fits in the left panel; gray points in the top plot show *p* obtained directly from simulations with *f* = 0. Solid lines show empirical fits. Right, validation of the empirical mapping. For each simulated dataset, *f* is estimated from its fitted *p* and known C and compared with the preset value used in the simulation. The estimation error, Δ*f* = *f*_estimated_ - *f*_preset_, is shown as a function of molecular concentration C (top, n = 41 in each) and preset *f* (bottom, n = 7 in each).

**Extended Data Fig. 3.**
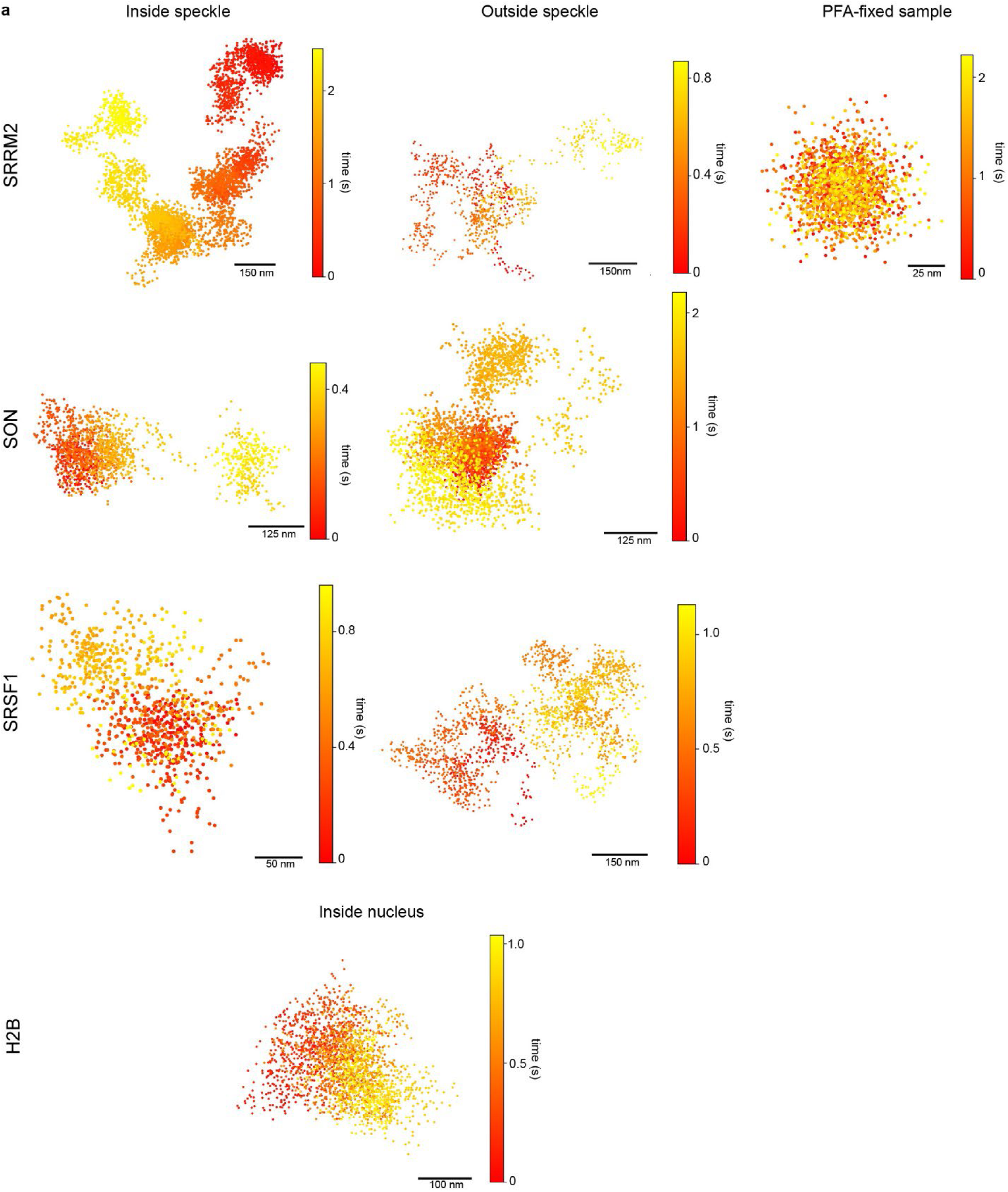
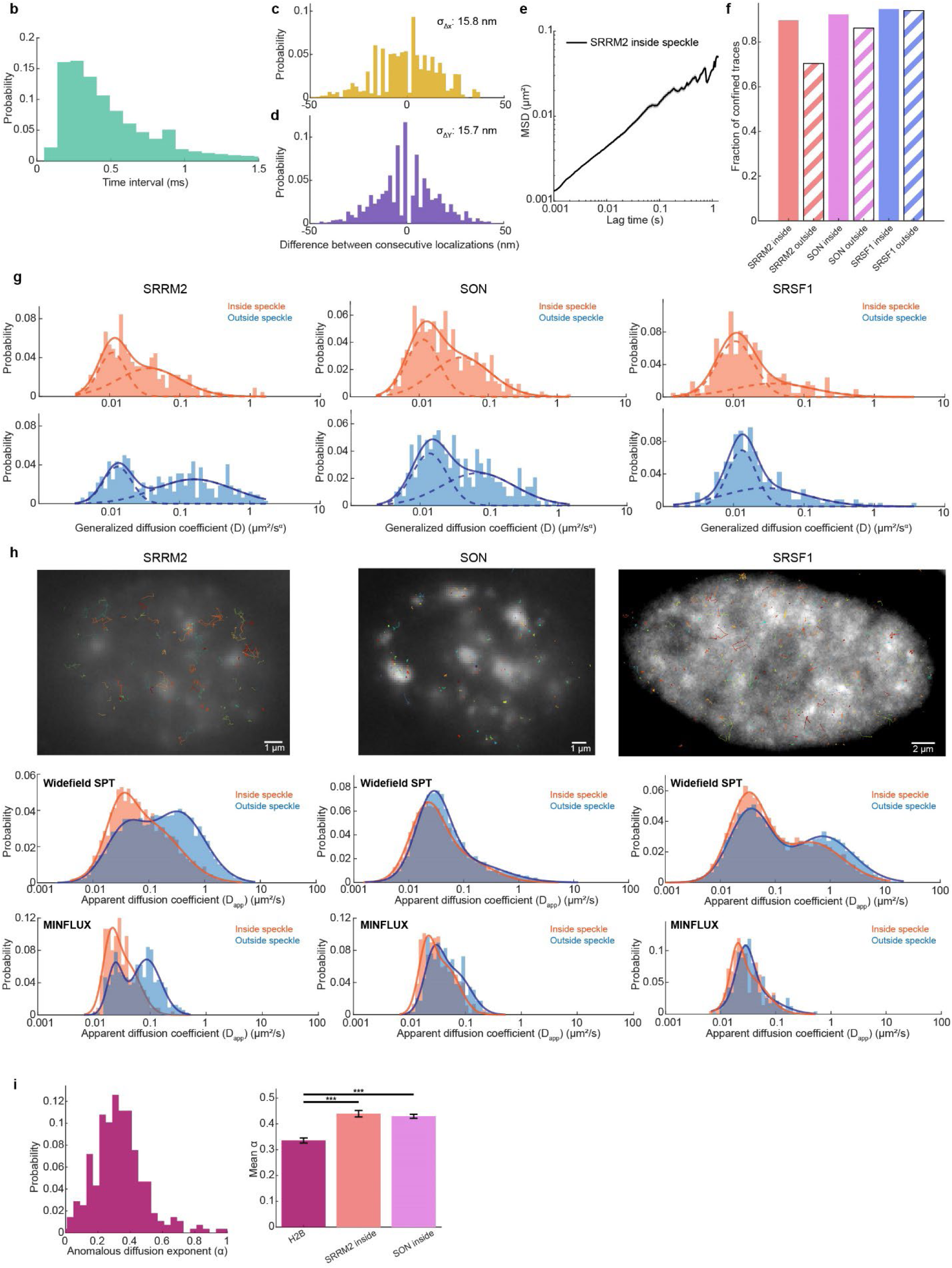
MINFLUX tracking reveals multi-state diffusion behaviors of speckle proteins. **(a)** Representative single-molecule trajectories of SRRM2, SON, SRSF1 inside (left) and outside (middle) nuclear speckles as well as a SRRM2 trajectory after PFA fixation (right). Bottom: representative single-molecule trajectories of H2B in U2OS cells. **(b)** Distribution of the temporal resolution of MINFLUX tracking, defined as the time interval between consecutive localizations of SRRM2 trajectories inside nuclear speckles. **(c, d)** Distributions of the displacements in the x (**c**) and y (**d**) dimensions between consecutive localizations of PFA-fixed SRRM2 trajectories. The standard deviation of the displacement distributions was used to estimate the localization precision. **(e)** Mean-squared displacement (MSD) as a function of lag time, averaged across full-length SRRM2 trajectories inside nuclear speckles. The shaded area represents SEM. **(f)** The fraction of confined trajectories (α < 0.7) for SRRM2 (red), SON (purple), and SRSF1 (blue) inside (solid) and outside (diagonal hatching) nuclear speckles. n = 225,321,524,290,209 and 236 trajectories, from left to right. **(g)** Distributions of generalized diffusion coefficients (D) for SRRM2 (left), SON (middle), and SRSF1 (right) inside (top, orange) and outside (bottom, blue) nuclear speckles. D values are estimated from MSD analyses of individual trajectories. The distributions are fitted to a sum of two Gaussians as a guide to the eye; dashed lines show the individual Gaussian components and solid lines show their sum. n = 225 and 321 trajectories for SRRM2, 524 and 290 trajectories for SON, and 209 and 236 trajectories for SRSF1, inside and outside speckles, respectively. **(h)** Widefield single-particle tracking (SPT) of speckle proteins SON (left), SRRM2 (middle), and SRSF1 (right). Top, representative widefield images of Halo-tagged proteins labeled with JF552, with trajectories shown in different colors. Distributions of apparent diffusion coefficients (D_app_) for each protein inside (orange) and outside (blue) nuclear speckles from widefield SPT (middle) and MINFLUX (bottom). D_app_ values are estimated from mean squared displacement (MSD) analysis of individual trajectories. The distributions are fitted to a sum of two Gaussians as a guide to the eye (solid lines). For MINFLUX: n = 225 and 321 trajectories for SRRM2, 524 and 290 trajectories for SON, and 209 and 236 trajectories for SRSF1, inside and outside speckles, respectively. For Widefield SPT, n = 2348 and 5534 trajectories for SRRM2, 7207 and 7192 trajectories for SON, and 5244 and 9669 trajectories for SRSF1, inside and outside speckles, respectively. Data are presented as mean ± SEM. (*) p ≤ 0.05, (**) p ≤ 0.01, and (***) p ≤ 0.001 (Mann–Whitney U test). **(i)** Left, distribution of the anomalous diffusion exponent (α) for Halo-H2B in U2OS. α values are obtained by fitting the MSD of individual trajectories to a power-law model. Right, mean α values for Halo-H2B (dark pink), SRRM2 (orange: inside speckle), and SON (purple: inside speckle). n = 278, 225, 524 trajectories, from left to right. Data are presented as mean ± SEM. (*) p ≤ 0.05, (**) p ≤ 0.01, and (***) p ≤ 0.001 (Mann–Whitney U test).

**Extended Data Fig. 4.**
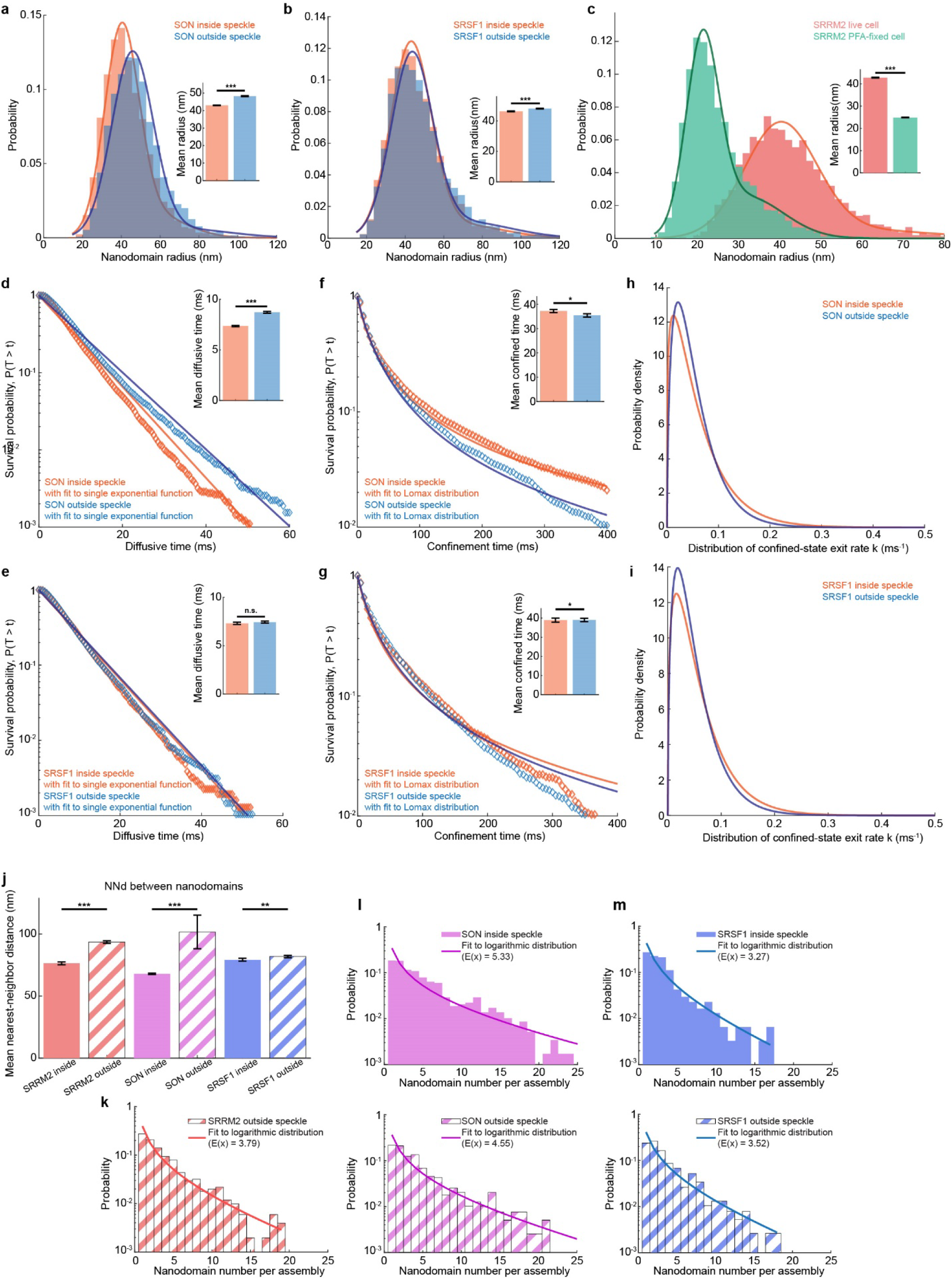
Speckle proteins transiently slow down within confined nanodomains. **(a, b)** Distributions of nanodomain radii for SON (**a**) and SRSF1 (**b**) inside (orange) and outside (blue) nuclear speckles (bars). Solid lines show two-Gaussian fits to the distributions as a guide to the eye. Inset represents mean nanodomain radii inside and outside speckles. n = 9609 and 3754 nanodomains for SON and 2,774 and 4,156 nanodomains for SRSF1, inside and outside speckles, respectively. **(c)** Distributions of nanodomain radii for SRRM2 inside nuclear speckles (red) and in PFA-fixed cells (green). Solid lines show two-Gaussian fits to the distributions as a guide to the eye. Inset shows mean nanodomain radii between live and fixed nuclei. n = 4070 nanodomains for SRRM2 live cell and 3992 nanodomains for SRRM2 PFA-fixed cell. **(d, e)** Distributions of diffusive-state dwell times for SON (**d**) and SRSF1 (**e**) inside (orange) and outside (blue) nuclear speckles. Diamond markers show the observed distributions, and solid lines show fits to a single-exponential function for dwell times >5 ms. Inset represents mean diffusive-state dwell times inside and outside speckles. n = 9060 and 5835 diffusive periods for SON and 3092 and 4777 diffusive periods for SRSF1, inside and outside speckles, respectively. **(f, g)** Distributions of confined-state dwell times for SON (**f**) and SRSF1 (**g**) inside (orange) and outside (blue) nuclear speckles. Diamond markers show the observed distributions, and solid lines show fits to a Lomax distribution. Inset represents mean confined-state dwell times inside and outside speckles. n = 9167 and 6016 confined periods for SON and 3267 and 5003 confined periods for SRSF1, inside and outside speckles, respectively. **(h, i)** Probability density functions of the confined-state exit rate k for SON (**h**) and SRSF1 (**i**) inside (orange) and outside (blue) nuclear speckles, obtained from the Lomax fits. Bar graphs in (**a**–**g**) show mean ± SEM. (*) p ≤ 0.05, (**) p ≤ 0.01, and (***) p ≤ 0.001 (Mann– Whitney U test). (j) The mean nearest-neighbor distance between adjacent nanodomains for SRRM2 (red), SON (purple), and SRSF1 (blue) inside (solid) and outside (diagonal hatching) nuclear speckles. Data are presented as mean ± SEM. (*) p ≤ 0.05, (**) p ≤ 0.01, and (***) p ≤ 0.001 (Mann–Whitney U test). n = 2156, 5101, 4977, 3682, 1997, and 3263, from left to right. **(k–m)** Distributions of the number of nanodomains per higher-order assembly for SRRM2 outside nuclear speckles (**k**) and for SRSF1 (**l**) and SON (**m**) inside (solid) and outside (diagonal hatching) nuclear speckles (bars), fitted to a logarithmic distribution (solid line). n = 506 assemblies for SRRM2 outside speckles; 306 and 381 assemblies for SRSF1, and 586 and 395 assemblies for SON, inside and outside speckles, respectively.

**Extended Data Fig. 5.**
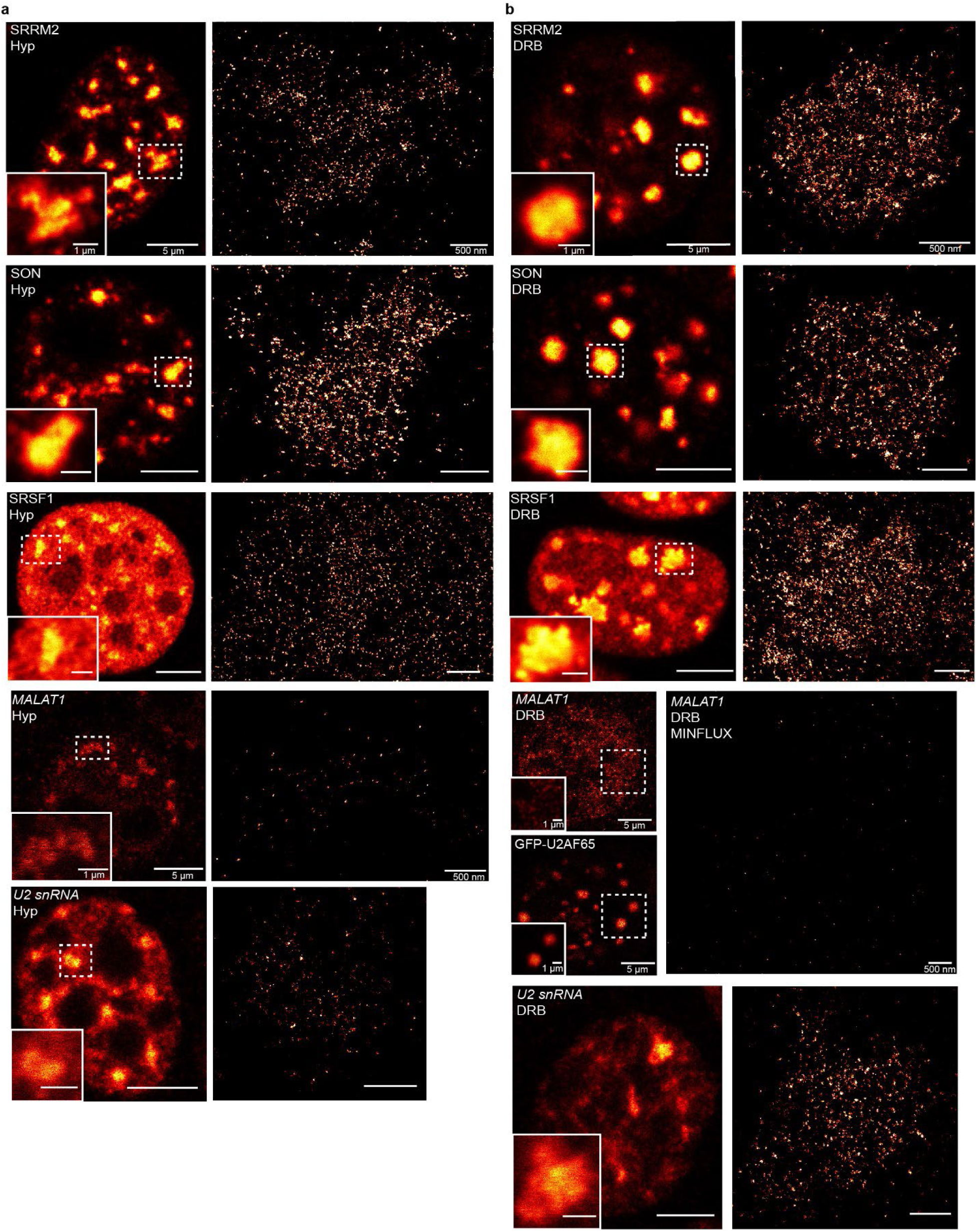
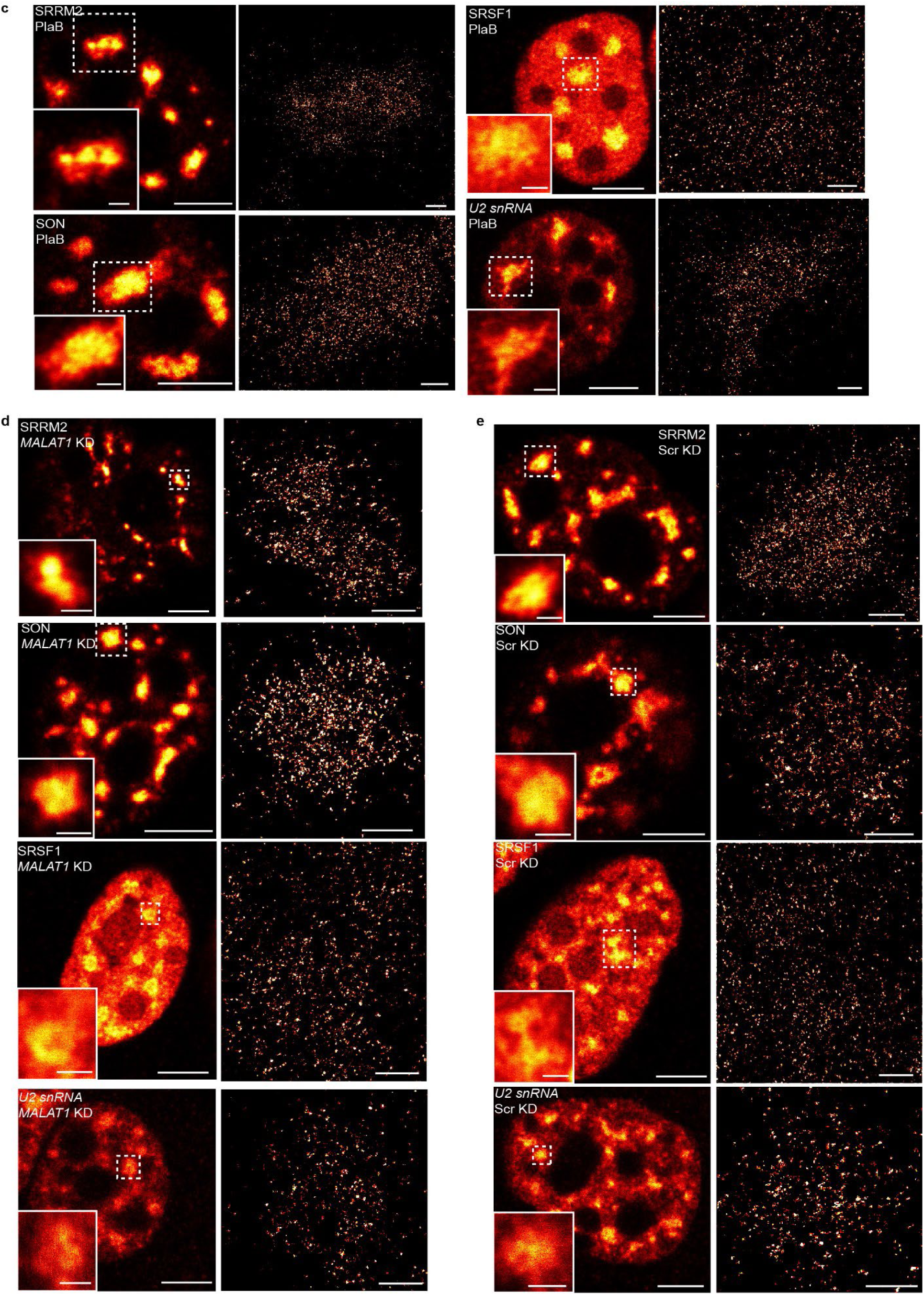
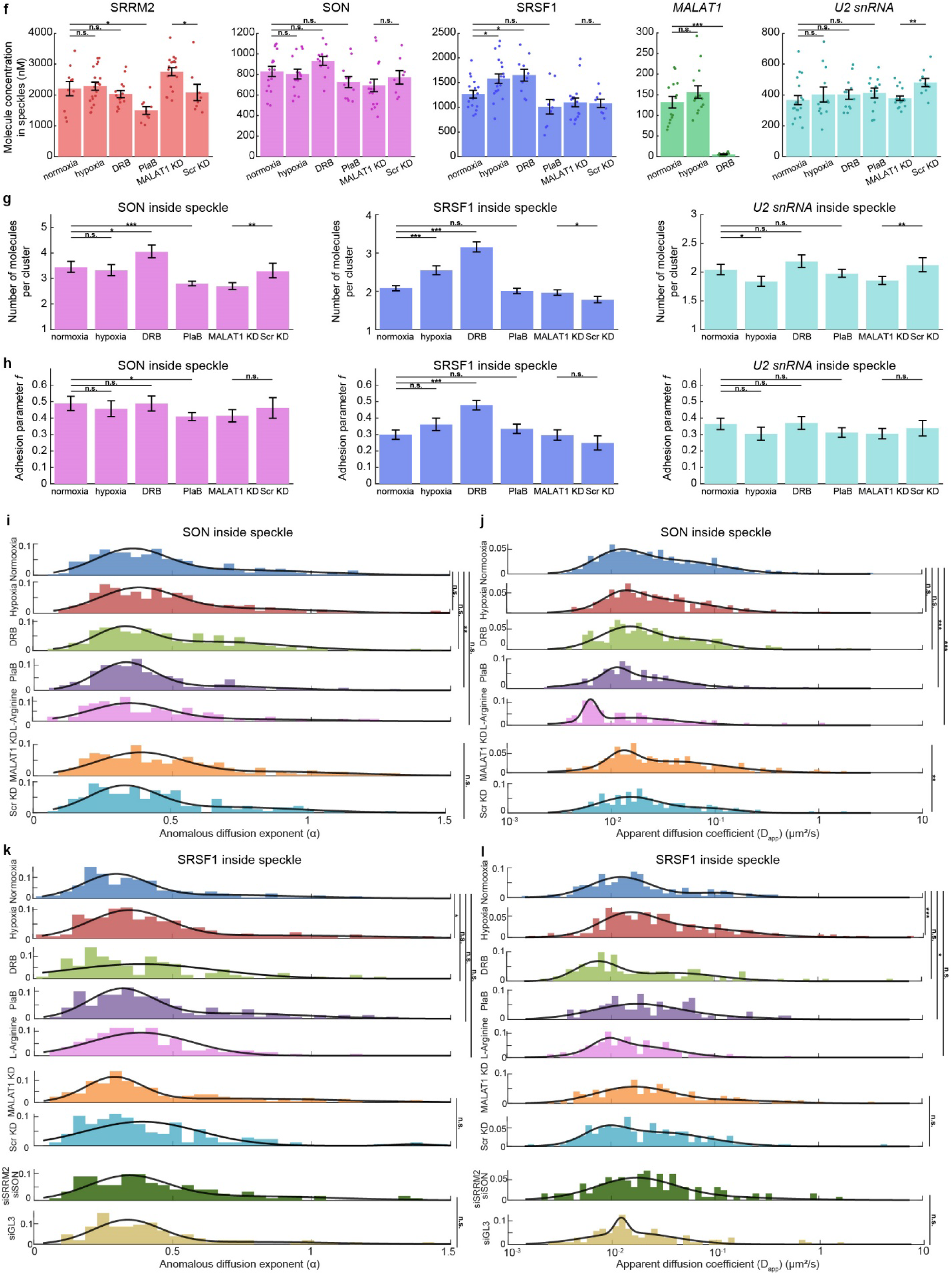
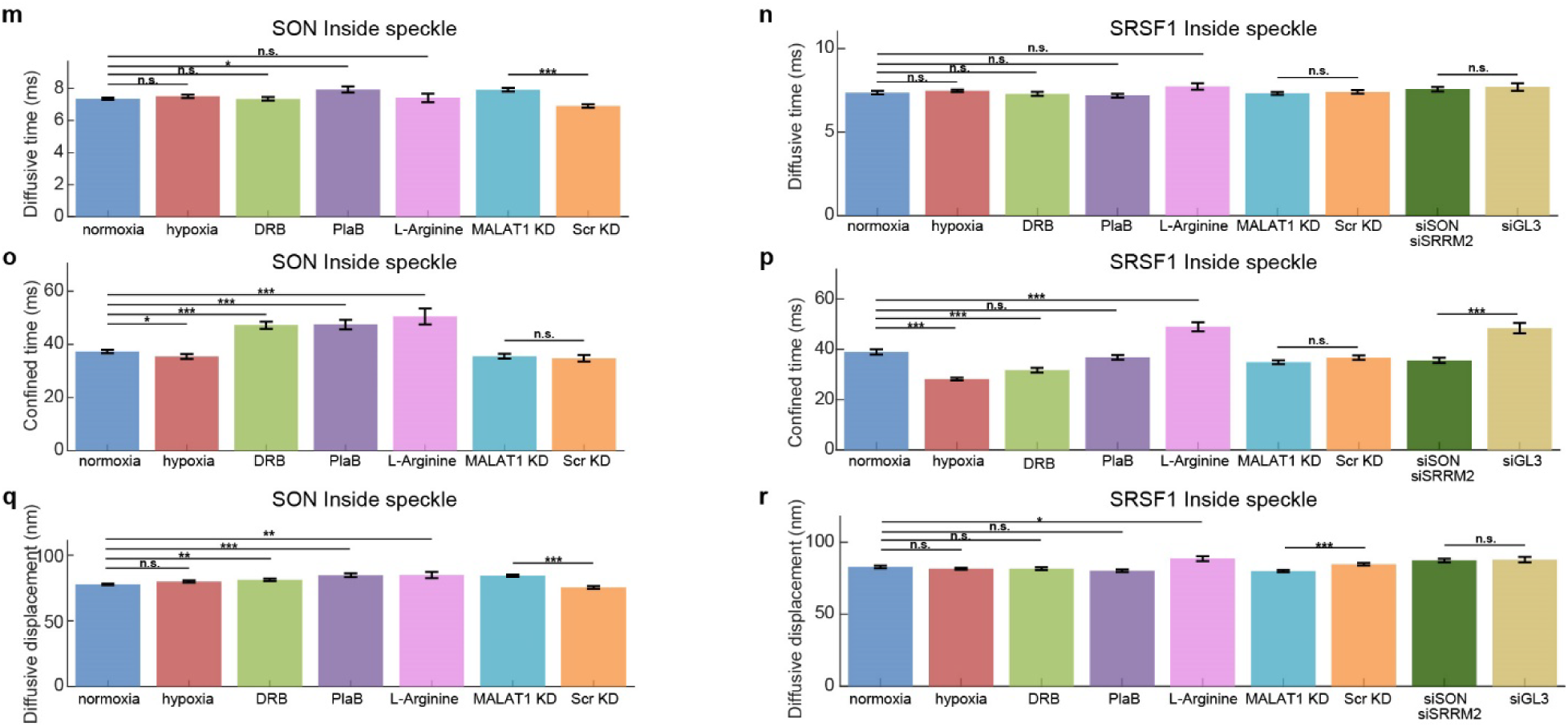
Perturbations alter the spatial organization and dynamics of speckle components. **(a–e)** Representative 2D MINFLUX images of SRRM2, SON, SRSF1, *MALAT1*, and *U2 snRNA* under hypoxia (**a**), DRB (**b**), PlaB (**c**), *MALAT1* knockdown (**d**), and scramble control oligo-treated cells (**e**). Left, confocal images acquired on the MINFLUX microscope showing the nucleus. Inset, magnified view of the selected nuclear speckle used for MINFLUX imaging. Right, 2D MINFLUX images of the corresponding nuclear speckles. **(f)** Concentrations of SRRM2, SON, SRSF1, *MALAT1*, and *U2 snRNA* inside nuclear speckles under different perturbations. Data are presented as mean ± SEM. (*) p ≤ 0.05, (**) p ≤ 0.01, and (***) p ≤ 0.001 (Mann–Whitney U test). From left to right, n = 10, 20, 14, 9, 18 and 8 speckles for SRRM2, n = 16, 14, 11, 11, 15 and 8 speckles for SON, n = 16, 19, 10, 9, 17 and 8 speckles for SRSF1, n = 14, 14 and 13 speckles for *MALAT1*, n = 19, 12, 12, 12, 17 and 10 speckles for *U2 snRNA*. **(g)** The expected number of molecules per cluster for SON (left), SRSF1 (middle), and *U2 snRNA* (right) inside nuclear speckles after different perturbations. **(h)** Adhesion parameter *f* for SON (left), SRSF1 (middle), and *U2 snRNA* (right) inside nuclear speckles after different perturbations. Bar graphs in (g and h) show means with 95% confidence intervals (CIs). Asterisks denote non-overlapping CIs: (*) 95%, (**) 99%, (***) 99.9%; n.s. denotes overlapping 95% CIs. **(i–j)** Distributions of the anomalous diffusion exponent (α) (**i**) and apparent diffusion coefficient (D_app_) (**j**) for SON inside nuclear speckles after different perturbations. The distributions are fitted to a sum of two Gaussians as a guide to the eye (solid lines). (*) p ≤ 0.05, (**) p ≤ 0.01, and (***) p ≤ 0.001 (Mann-Whitney U test). n = 524, 275, 343, 233, 102, 260, and 209 trajectories, from top to bottom. **(k–l)** Distributions of the anomalous diffusion exponent (α) (**k**) and apparent diffusion coefficient (D_app_) (**l**) for SRSF1 inside nuclear speckles after different perturbations. The distributions are fitted to a sum of two Gaussians as a guide to the eye (solid lines). (*) p ≤ 0.05, (**) p ≤ 0.01, and (***) p ≤ 0.001 (Mann-Whitney U test). n = 209, 292, 129, 157, 146, 210, 187, 123, and 110 trajectories, from top to bottom. **(m–n)** The mean diffusive-state dwell times of SON (**m**) and SRSF1 (**n**) after indicated perturbations. **(o–p)** The mean confined-state dwell times of SON (**o**) and SRSF1 (**p**) after indicated perturbations. **(q–r)** The mean diffusive displacement of SON (**q**) and SRSF1 (**r**) after indicated perturbations. Bar graphs in (m–r) show mean ± SEM. (*) p ≤ 0.05, (**) p ≤ 0.01, and (***) p ≤ 0.001 (Mann– Whitney U test). For SON, n = 9060, 3922, 3037, 1456, 511, 4577, and 2388 diffusive periods in (**m, q**), and 9167, 3987, 3155, 1526, 555, 4772, and 2466 confined periods in (**o**), from left to right. For SRSF1, n = 3092, 8685, 2896, 3018, 1477, 4964, 3673, 2577, and 1159 diffusive segments in (**n, r**), and 3267, 8887, 2974, 3159, 1570, 5123, 3854, 2724, and 1223 confined segments in (**p**), from left to right.

## Notes

### Competing Interest Statement

Susan M. Freier is an employee of Ionis inc, and receives salary from the company.

